# Impact of a transient neonatal visual deprivation on the development of the ventral occipito-temporal cortex in humans

**DOI:** 10.1101/2024.11.30.625697

**Authors:** Stefania Mattioni, Mohamed Rezk, Xiaoqing Gao, Junghyun Nam, Zhong-Xu Liu, Remi Gau, Valérie Goffaux, Andrea I. Constantino, Hans Op de Beeck, Terri Lewis, Daphne Maurer, Olivier Collignon

**Author notes:** Correspondence should be addressed to: Stefania Mattioni and Olivier Collignon.

## Abstract

How does experience shape the development of visual brain regions? We demonstrate that a transient period of visual deprivation early in life in humans leads to permanent alteration in the function of the early visual cortex (EVC), while leaving the categorical coding of downstream ventral occipito-temporal cortex (VOTC) mostly unaffected. We used fMRI to characterize the brain response to five categories (faces, bodies, objects, buildings and words) in a rare group of cataract-reversal individuals who experienced a transient period of blindness early in life, and in matched participants with typical visual development. We show that the encoding of low-level visual properties is impaired in EVC in cataract-reversal participants, while the categorical response in the VOTC is preserved. When altering the visual properties of our stimuli to mimic in controls the deficit of EVC response of the cataract, we observe a cascading alteration of the categorical coding from EVC to VOTC that is not observed in the cataract-reversal group. Our results show that a typical visual categorical organization develops in VOTC despite a period of visual deprivation early in life and in the presence of impaired processing in EVC. A deep neural network exposed to a visual diet that emulates the experience of our participants reproduced our brain observations within an artificial system. These findings emphasize how specific visual regions are differently affected by a period of visual deprivation early in life, with information encoded in EVC being permanently affected, while the categorical coding in VOTC shows resilience to deprivation.

## Introduction

The study of the consequence of an early and transient period of visual deprivation has long served as a compelling experimental model to causally test the need of sensory experiences for the functional development of brain regions (Hubel & Wiesel, 1963).

Seminal studies have unveiled the existence of sensitive and critical developmental periods during which early life experience significantly shapes the functional tuning of the brain (Knudsen, 2004). Studies in mice (Gordon & Stryker, 1996; Prusky & Douglas, 2003), cats (Hubel & Wiesel, 1963; Wiesel & Hubel, 1963a, 1963b, 1965) and monkeys (Le Vay et al., 1980) have demonstrated profound and enduring effects of a brief and transient period of monocular or binocular deprivation in early life on responses of the early visual cortex (EVC) (for comprehensive reviews, see Mitchell & Maurer, 2022). Similarly, humans who experience transient postnatal blindness due to dense bilateral cataracts show permanently reduced visual acuity (Lewis et al., 1995; Lewis & Maurer, 2009; Sinha & Held, 2012; Zohary et al., 2022) and alteration in the response and retinotopic organization of the EVC (Heitmann et al., 2023; Sourav et al., 2018).

The impact of a transient neonatal visual deprivation beyond EVC has been less explored; yet it is commonly assumed that these regions are altered due to their connection to, and dependence from, lower-level properties extracted from EVC (M. J. Arcaro & Livingstone, 2021; M. Arcaro & Livingstone, 2024). It has, for instance, been suggested that the transient absence of early visual experience could disrupt the typical development of category selectivity in the ventral occipito-temporal cortex (VOTC), potentially even to a greater extent than it impacts EVC (M. J. Arcaro & Livingstone, 2021; M. Arcaro & Livingstone, 2024; Sourav et al., 2018, 2020). Most of the studies that have explored categorical representations in VOTC among cataract-reversal individuals have focused on the processing of faces. Some of these studies have reported impairments in tasks involving the localization of internal facial features (De Heering & Maurer, 2014; Le Grand et al., 2001; Robbins et al., 2010), face memory tasks (De Heering & Maurer, 2014), ratings of facial attractiveness (Gupta et al., 2023), automatic gaze following (Zohary et al., 2022) accompanied by abnormal (Mondloch et al., 2013) or absent (Röder et al., 2013) electrophysiological responses (N170) to face stimuli and altered connectivity within the face-processing network (Grady et al., 2014). Similarly, a study involving monkeys raised without selective exposure to faces has reported the absence of face domain in the brain after sight recovery (M. J. Arcaro & Livingstone, 2017).

These results are however not without controversies. Other studies showed that cataract-reversal individuals can successfully perform several tasks including face detection (Mondloch et al., 2013), face/nonface categorization (Gandhi et al., 2017), discrimination among individual faces based on the shape of the eyes, the mouth or the external contour (Mondloch et al., 2010) or the extraction of social information from faces, such as facial and emotional expression (Gao et al., 2013; Gilad-Gutnick et al., 2024). In the same line, monkeys raised without exposure to any faces exhibited a preference for human and monkey faces in photographs, along with the ability to discriminate between human faces as well as monkey faces (Sugita, 2009).

Remarkably, no previous studies have simultaneously investigated the concurrent development of early and higher-level visual regions across different visual categories in the context of cataract-reversals. Moreover, so far, most studies comparing brain responses between cataract-reversal and controls have relied on univariate techniques. However, univariate and contrast approaches applied to a restricted set of stimuli are limited in their capacity to reveal the precise nature of the underlying brain computations that may differ between cataracts and controls, and how these potential changes in computation vary according to brain regions (Haxby et al., 2014).

Here, we examined functional differences along the ventral occipito-temporal stream between cataract reversal individuals and controls using decoding and representational similarity analyses (RSAs) (Kriegeskorte, 2008; Mattioni, 2024) applied to functional Magnetic Resonance Imaging (fMRI) signals and artificial neural network models. Using a diverse set of images belonging to multiple visual categories including faces, bodies, houses, tools and words, we concomitantly probed alteration in the brain coding of either low-level (e.g. spatial frequencies) or high-level (e.g. categorical) visual features of afferent visual input. We also included a control fMRI experiment where we presented modified visual stimuli to additional control subjects to simulate the visual deficits associated with cataract reversal (i.e.., reduced visual acuity and nystagmus), allowing us to account for partially impaired vision at the time of testing. Additionally, we directly compared the brain’s response along the visual cortical hierarchy with the representation of different deep neural network (DNN) layers, thought to emulate how information is sequentially extracted in VOTC (Cichy et al., 2016).

We observed that despite impaired encoding of low-level visual properties in EVC among cataract-reversal participants, a typical categorical organization develops in VOTC. We reproduced such observation in an artificial neural network fed with a developmental visual diet mimicking the one of our participants.

## Materials and methods

### Participants

Fifteen cataract-reversal patients (7 males, mean age ± SD= 28.26 ± 6.32 years, age range= 18 - 39 years, see Table 1) and seventeen matched controls (11 males, mean age ± SD= 29.12 ± 5.71 years, age range= 21 – 39 years) participated in the experiment. An additional group of fourteen matched control subjects (7 males, mean age ± SD= 27.86 ± 4.91 years, age range= 21 – 39 years) participated in a control experiment in which the visual properties of the stimuli were altered. All participants were naive to the purpose of the experiment. The protocol was approved by the research ethics committee of the University of Toronto and McMaster University. A written consent was acquired from all participants. Participants were monetarily compensated for their participation. Prior to testing, participants underwent a training session where they were familiarized with the experiments and the required tasks. Based on the poor performance in the task during the fMRI data acquisition, we excluded one subject from the control group, one subject from the cataract-reversal group and one subject from the control experiment; see the section on behavioral results in the fMRI for further details.

**Table 1.** Characteristics of reversal cataract participants.

| Subjects | Age(y) | Sex | Length of deprivation (days) | Acuity (LogMar) | Nistagmus |
| --- | --- | --- | --- | --- | --- |
| CAT1 | 39 | M | 124 | 0.17 | YES |
| CAT2 | 34 | F | 47 | 0.14 | YES |
| CAT3 | 22 | F | 84 | 0.28 | YES |
| CAT4 | 27 | F | 132 | 0.20 | N.A. |
| CAT5 | 38 | F | 44 | 0.19 | N.A. |
| CAT6* | 27 | F | 107 | 0.31 | YES |
| CAT7 | 23 | F | 6 | 0.25 | YES |
| CAT8 | 32 | M | 94 | 0.61 | YES |
| CAT9 | 26 | M | 77 | 0.52 | YES |
| CAT10 | 37 | M | 143 | 0.09 | YES (mild) |
| CAT11 | 18 | F | 10 | 0.14 | NO |
| CAT12 | 25 | F | 65 | 0.05 | YES (mild) |
| CAT13 | 26 | M | 21 | 0.26 | YES |
| CAT14 | 27 | M | 118 | 0.09 | NO |
| CAT15 | 23 | M | 3 | 0.04 | NO |

*\*Sub CAT6 has been excluded based on the low performance in the fMRI task.*

### Assessment of visual acuity

The visual acuity was acquired for the cataract-reversal (see Table 1) and the control subjects undergoing the main experiment using the Freiburg Visual Acuity Test (Bach, 1996, 2006). Participants were tested at a distance of 210 cm from the stimulation monitor (CRT monitor of screen width of 53.5 cm, and screen resolution of 800 x 600 pixels). The test run comprised 30 Landolt-C trials that were presented in a random orientation. The size of the stimuli was adjusted on each trial based on the previous responses to estimate the most probable visual acuity threshold by using a maximum-likelihood staircase procedure (Lieberman & Pentland, 1982). The participants performed a forced choice task where they indicated the orientation of the C-stimulus gap (up, down, right, left). The Logarithm of the Minimum Angle of Resolution (logMar) was used as a measure of visual acuity. During this visual acuity test, the subjects wore the same corrective glasses they used in the scanner during the fMRI experiment. Therefore, the visual acuity measured represents a corrected level, reflecting the maximal acuity the subjects could achieve at the time of the test. The visual acuity of the two groups were compared using a two-sample t-test. Furthermore, the visual acuity values were used to conduct correlations with brain data.

### Stimuli and Procedure for fMRI experiment

Visual stimuli were projected on a screen (frame rate: 60 Hz; screen resolution 1920 x 1080 pixels) behind the scanner. Participants viewed the screen (distance from head= 45 cm) through a mirror mounted on the MRI head coil. Participants performed a task during the experiment by responding with two MR-compatible response buttons.

The stimulus set included 6 images in each of 5 different categories: bodies, faces, houses, tools and words (30 images; see fig. 1A). The images were black and white pictures (500X500 pixels) collected from internet. They were placed in the center of the screen on a gray (129 RGB) background.

**Figure 1.**
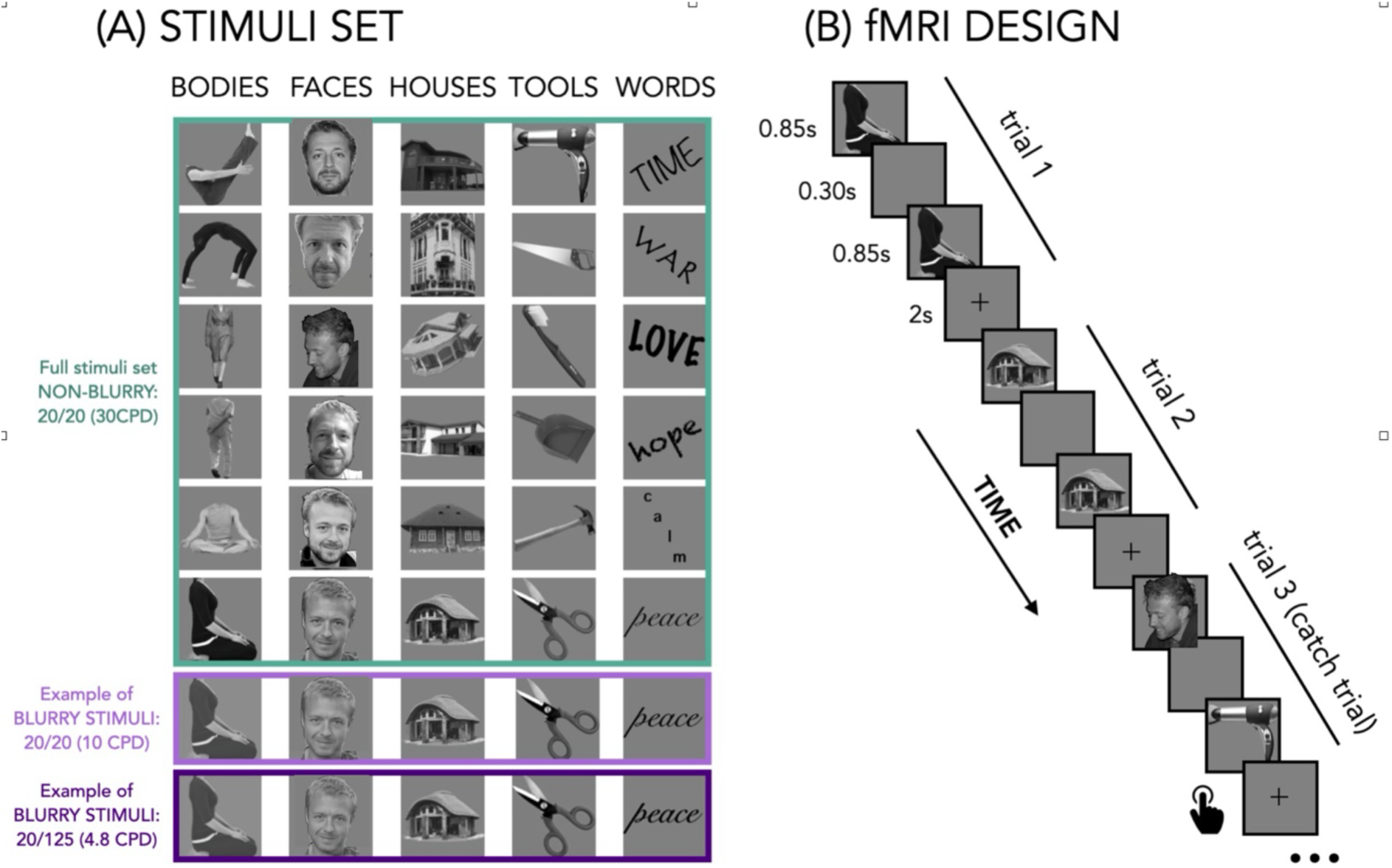
Stimuli Set and Experimental Design. **Panel A** displays the stimuli used in the experiment. The green box encompasses the full stimuli set used in the main (not-blurry) experiment, with each column representing a category and each row depicting different exemplars within the category, totaling 30 images. Pictures of the “FACE” category are all from the last author for copyright reason and therefore do not represent the faces presented in the experiments that were all from different actors. Additionally, the light and dark purple boxes illustrate one exemplar per category in two control experiments where the images were spatially filtered at two different levels of blurriness: 10 Cycles per degree (CPD), representing the average visual acuity level of the reversal-cataract group (logMar=0.21), and 4.8 CPD, which denotes the lowest level of acuity recorded among the cataract-reversal individuals (logMar=0.61). All the stimuli, without any compression, can be found in the OSF project (DOI 10.17605/OSF.IO/BECDR) of this study at this [link](https://osf.io/2cte7/?view_only=3a96c24e49034621b22c8ef2cd258530). **Panel B** depicts the experimental design during fMRI.

Before entering the scanner, each participant was familiarized with the stimuli to ensure perfect recognition. In the fMRI event-related experiment each trial consisted of the same stimulus repeated twice. Rarely (10% of the occurrences), a trial was made up of two different consecutive stimuli (catch trials). Only in this case, participants were asked to press a key with the right index finger if the second stimulus belonged to the living category and with their right middle finger if the second stimulus belonged to the non-living category. This procedure ensured that the participants attended and processed the stimuli. Each pair of stimuli lasted 2s (850 msec per stimulus interleaved with 300 msec of a blank screen) and the inter-trial interval (i.e. fixation cross) was 2s long for a total of 4s for each trial (see fig.1B). Within the fMRI session participants underwent 5 runs, with the exception of one control subject that, due to technical issues, underwent only 4 runs. Each run contained 2 repetitions of each of the 30 stimuli, 6 catch trials and two 20s-long fixation periods without stimuli (one in the middle and another at the end of the run). The total duration of each run was 304s. The presentation of trials was pseudo-randomized: two stimuli from the same category (i.e. bodies, faces, houses, tools and words) were never presented in subsequent trials. The stimuli delivery was controlled using Matlab R2016b (https://www.mathworks.com) Psychophysics toolbox (http://psychtoolbox.org).

In the control experiment that we ran in a separate group of participants, the visual properties of the images were altered to mimic the lower acuity and nystagmus of cataract-reversal subjects. Aside from image alteration, the procedure was identical to the one described for the original experiment. Nystagmus is a condition characterized by involuntary, rhythmic eye movements that can occur in people who have been treated for bilateral congenital dense cataract (10 of our cataract-reversals participants had a nystagmus, see Table 1). To simulate the nystagmus of cataract-reversal participants we applied to the visual stimulation a pendular movement in the horizontal plane. The like-nystagmus movement was applied to the image for the total time of the presentation (i.e. 850 msec) with a frequency of 3.5 hz. The maximal displacement of each image was of 150 pixels, corresponding to 3.23 degrees of visual angle (corresponding to the larger gaze’s displacement among the cataract subjects, see fig. 2C). Therefore, each image was shifting of 3.23 degrees in the horizontal direction for 3 times during the 850msec of presentation. To mimic the lower acuity of the cataract-reversals group we blurred the original stimulus set, applying a lowpass filter. We selected 2 different cutoff levels: 10 cycles per degree (CPD) corresponding to a visual acuity of 20/60 (average acuity level of our cataract-reversal group corresponding to a logMar of 0.21) and 4.8 CPD corresponding to an acuity of 20/125 (lower acuity level of the cataract-reversal group corresponding to a LogMar of 0.61). Therefore, we created two versions of the control experiment. In both versions the nystagmus was applied in the same way, while the level of blurring was either 10 CPD (CON-Blurry 1) or 4.8 CPD (CON-Blurry 2). All these stimuli, without any compression, can be found in the OSF project (DOI 10.17605/OSF.IO/BECDR) at this [link](https://osf.io/2cte7/?view_only=3a96c24e49034621b22c8ef2cd258530). A separate group of 14 subjects took part in both versions of the control experiment. Each subject performed 5 runs of the control experiment-blurry 1 and 5 runs off the control experiment-blurry 2.

**Figure 2.**
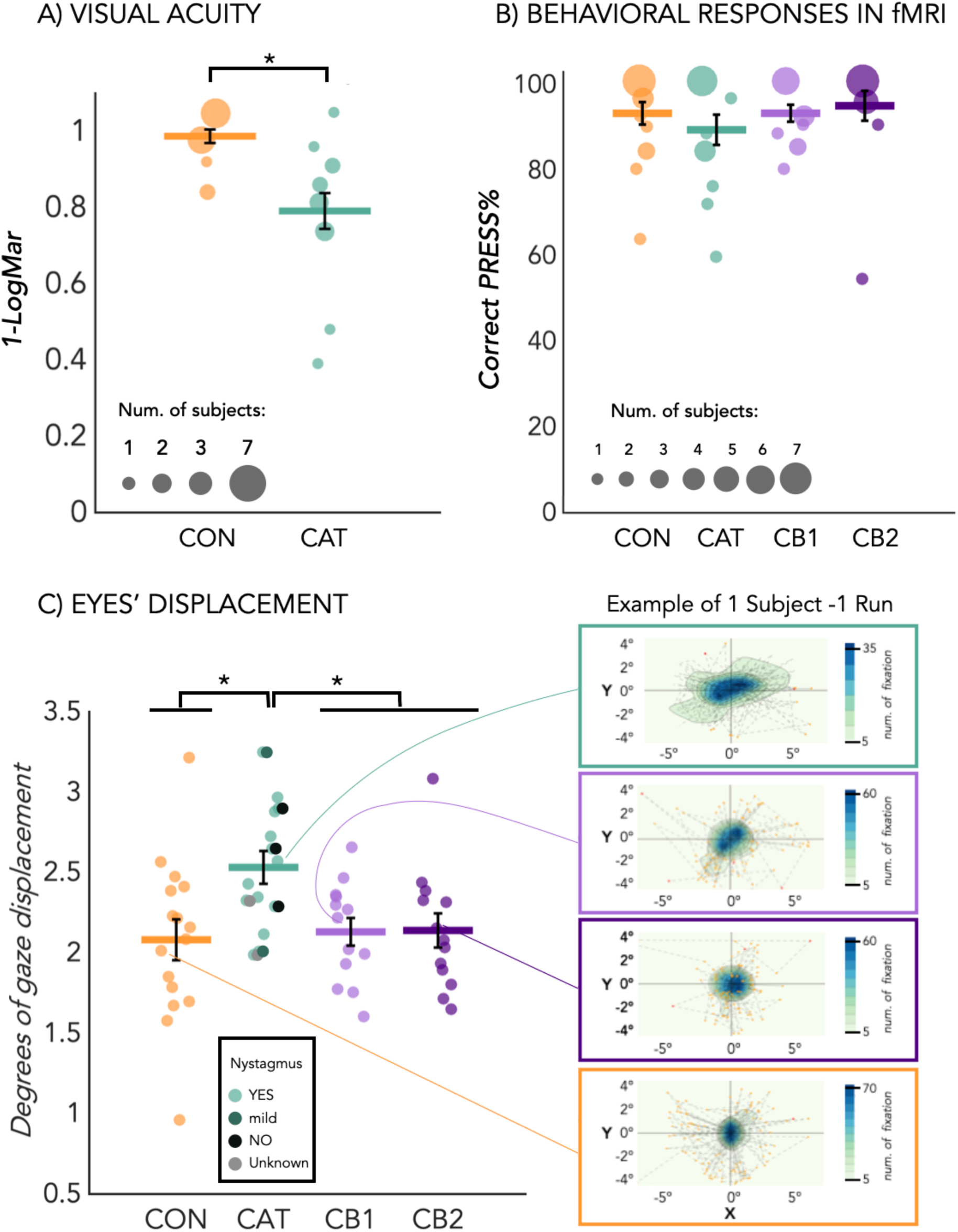
Visual Acuity, Behavioral Response, and Eye Displacement Across Experimental Conditions. **Panel A:** Visual acuity, computed as 1 minus the Logarithm of the Minimum Angle of Resolution (logMAR), is depicted for Cataract Reversals (CAT) and Control (CON) subjects. Dot size reflects subject count, with smaller dots representing one subject and larger dots indicating up to seven subjects. Horizontal lines denote group means, and black bars signify standard errors. **Panel B:** Behavioral response during fMRI is shown for CAT, CON, Controls Blurry-1 (CB1), and Controls Blurry-2 (CB2) groups. Dot sizes correspond to subject counts, with red-circled dots indicating outliers (< 2 standard deviations from the group mean). Mean values and standard errors are computed after excluding outliers. **Panel C:** Eye displacement during the fMRI experiment, analyzed using deepMReye, is compared across CAT, CON, CB1, and CB2 groups. Each dot represents one subject, with horizontal bars indicating group averages and vertical black lines denoting standard errors. Additionally, on the right side, an example of the deepMReye output for one average subject in each group during one run of the experiment is provided. Movement intensity is represented by color spread, with darker shades indicating more frequent fixation.

### MRI data acquisition

Structural and functional data were acquired at the Sherman Health Science Research Centre (York University, Toronto, Canada) using a Siemens MAGNETOM 3T PrismaFit MRI scanner with a standard 64 channel head coil. Structural high-resolution T1-weighted MPRAGE scans were acquired using parallel imaging (GRAPPA factor = 2). The acquisition parameters for the T1 images included a repetition time (TR) of 2300 ms, echo time (TE) = 2.62 ms, voxel size = 1 mm isotropic voxels, number of slices=192, and a flip angle of 9°. Functional scans were acquired using T2*-weighted echoplanar BOLD imaging. The acquisition parameters included simultaneous interleaved multi-slice acquisition using parallel imaging with multiband acceleration = 2, phase encoding acceleration = 3, TR = 1170 ms, TE = 30ms, voxel-size = 2 mm isotropic, number of slices = 51, flip angle = 66°, and echo-spacing 0.68 ms. The first four initial scans of each run were discarded to allow for equilibrium magnetization.

### fMRI preprocessing

fMRI data was preprocessed in statistical parametric mapping software (SPM12 – Wellcome Department of Imaging Neuroscience, University College London, UK) implemented in Matlab R2016b (Mathworks, Inc.). Preprocessing steps included slice time correction, EPI alignment to the mean functional image with a 2nd degree B-spline interpolation, co-registration of the functional volumes to the structural image and normalization to the Montreal Neurological Institute (MNI) template. For the univariate analysis only, we also performed a spatial smoothing with a Gaussian kernel of FWHM of six millimetres on the volume time series.

### Eye movement extraction from fMRI data

Ten out of our 16 cataract-reversal participants presented nystagmus. To control for the possible different pattern of eye movement in our groups, we used DeepMReye, an open source framework for eye-tracking that does not require a camera. It is based on a convolutional neural network (CNN) that reconstructs the viewing behavior of individuals directly from the MR signal of their eyeballs (Frey et al., 2021). At each image acquisition, DeepMReye captures the multi-voxel pattern of the eyes and use a CNN to predict the gaze location based on that pattern. To train the CNN, the researchers required an independent measure of gaze location, which was obtained from previous studies using camera-based eye tracking or fixation targets (see below for further details). Notably, although independent gaze information was necessary for the CNN training, it is not required when applying the trained CNN to new data (Krajbich, 2021).

To obtain the “eye-tracking” results from our fMRI data, we performed the preprocessing using bidsMReye (version 0.3.0+24.gbe1f5da.dirty), a BIDS app relying on deepMReye (@deepmreye) to decode eye motion from fMRI time series data.

The data of each BOLD run underwent co-registration conducted using Advanced Normalization Tools (ANTs, RRID:SCR_004757) within Python (ANTsPy). First, each participant’s mean EPI was non-linearly co-registered to an average template. Second, all voxels within a bounding box that included the eyes were co-registered to a preselected bounding box in our group template to further improve the fit.

Each voxel within those bounding box underwent two normalization steps. First, the across-run median signal intensity was subtracted from each voxel and sample and was divided by the median absolute deviation over time (temporal normalization). Second, for each sample, the mean across all voxels within the eye masks was subtracted and divided by the standard deviation across voxels (spatial normalization). Importantly, this method can decode gaze position at a temporal resolution higher than the one of the imaging protocol (sub-TR resolution) (Frey et al., 2021).

Voxels time series were used as inputs for generalization decoding using a pre-trained model 1 to 6 from deepMReye from OSF. This model was trained on the following datasets: guided fixations (@alexander_open_2017), smooth pursuit (@nau_real-motion_2018, @polti_rapid_2022, @nau_hexadirectional_2018), free viewing (@julian_human_2018).

For each run the following values were computed: (1) the variance for the X gaze position; (2) the variance for the Y gaze position; (3) the framewise gaze displacement; (4) the number of outliers for the X gaze position; (5) the number of outliers for the Y gaze position; (6) the number of outliers for the gaze displacement. Outliers were robustly estimated using an implementation of @carling_resistant_2000.

We used the average amount of gaze displacement across runs, after excluding the outliers, to test with independent samples T-tests whether there was a difference in the amount of eye movement between our groups (fig. 2C).

We included the variance for the X and Y gaze positions as regressors of no-interest in our GLM (see next paragraph), to control for the impact of eye movement on the fMRI data activity.

#### General linear model

The pre-processed images for each participant were first analyzed using a general linear model (GLM).

For the univariate analyses, for each subject, a design matrix was formed using a predictor for each stimulus category (bodies, faces, houses, tools, words) in each run. These regressors of non-interest were also added: 1 regressor of no-interest for the catch trials, 6 head-motion regressors of no-interest, 2 eye movement regressors of no-interest (the variance for the X gaze position and the variance for the Y gaze position) and 1 constant.

For the multivariate analyses, for each of the 5 runs we included 40 regressors: 30 regressors of interest (each stimulus), 1 regressor of no-interest for the catch trials, 6 head-motion regressors of no-interest, 2 eye movement regressors of no-interest (the variance for the X gaze position and the variance for the Y gaze position) and 1 constant. From the GLM analysis we obtained a β-image for each stimulus (i.e. 30 pictures) in each run, for a total of 150 (30 x 5) beta maps.

#### Statistical procedure and brain masks for TFCE correction

For each one-sample t-test within-group and two-sample t-tests performed to compare the effects between groups, results were corrected using the non-parametric threshold free cluster enhancement (TFCE) method combined with a FWE correction (Smith & Nichols, 2009).

All univariate and searchlight analyses in the paper were run whole brain. But the threshold free cluster enhancement (TFCE) correction for the two-sample t-tests was applied within specific masks, based on a-priori hypotheses. We were a priori interested in how our visual stimuli were processed for their low-level visual features and for their high-level categorical visual properties. We therefore created a mask including the primary visual cortex (V1) and the ventral occipito-temporal cortex (VOTC) by combining these regions from the JuBrain Anatomy Toolbox bilaterally (a.k.a. SPM Anatomy Toolbox - v2.2b, (Eickhoff et al., 2005): the *Human occipital cytoarchitectonic area 1, hOc1(Amunts et al., 2000)*, *Area FG1* situated in the posterior and medial regions of the fusiform gyrus, *Area FG2* located in the posterior fusiform gyrus and extending to the occipito-temporal sulcus, *Area FG3*, anterior to FG1, placed in the medial fusiform gyrus and expanding till the collateral sulcus (Lorenz et al., 2015) and *Area FG4,* anterior to FG2, including the lateral portion of the fusiform gyrus (FG) and extending to the occipito-temporal sulcus (Lorenz et al., 2015). For a similar parcellation of the human ventral visual stream see also (Rosenke et al., 2018). These masks can be found at this [link](https://osf.io/vm7p2/?view_only=875597cd92184e2b81b2b96ee65db37c), on the OSF project (DOI 10.17605/OSF.IO/BECDR) linked to this work.

The correction was applied either across the full mask, including bilateral V1 and bilateral VOTC (for univariate analysis [all categories > baseline] and split-half analysis), solely in the VOTC mask (for univariate analysis [each category > other categories], for RSA with categorical model and 5-way categorical decoding), or exclusively in the V1 mask (for RSA with Hmax-C1 model and within-category decoding), based on a priori hypotheses that are explained in each respective section below.

#### Univariate analysis

Using SPM12 we computed within and between groups contrast maps comparing (1) all visual stimuli vs baseline; (2) bodies vs all other categories; (3) faces vs all other categories; (4) houses vs all other categories; (5) tools vs all other categories; (6) words vs all other categories. Since the last contrast did not show any activation across groups, even at a low correction threshold, we also ran and reported the contrast *words > houses*. These contrast images were further spatially smoothed with a Gaussian kernel of 6mm FWHM before conducting group-level analyses. They were input into a series of corresponding one-sample t-tests within each of the four groups and two-sample t-tests between each possible group comparison (see fig.3 and SI fig.1). For the groups’ contrasts (two-sample t-tests) the TFCE correction was applied within the general visual mask including both V1 and VOTC (for details about the masks, see the section above: *Statistical procedure and brain masks for TFCE correction)*, since for this analysis the difference in brain activity between groups could emerge for both visual low-level and/or categorical features.

**Figure 3:**
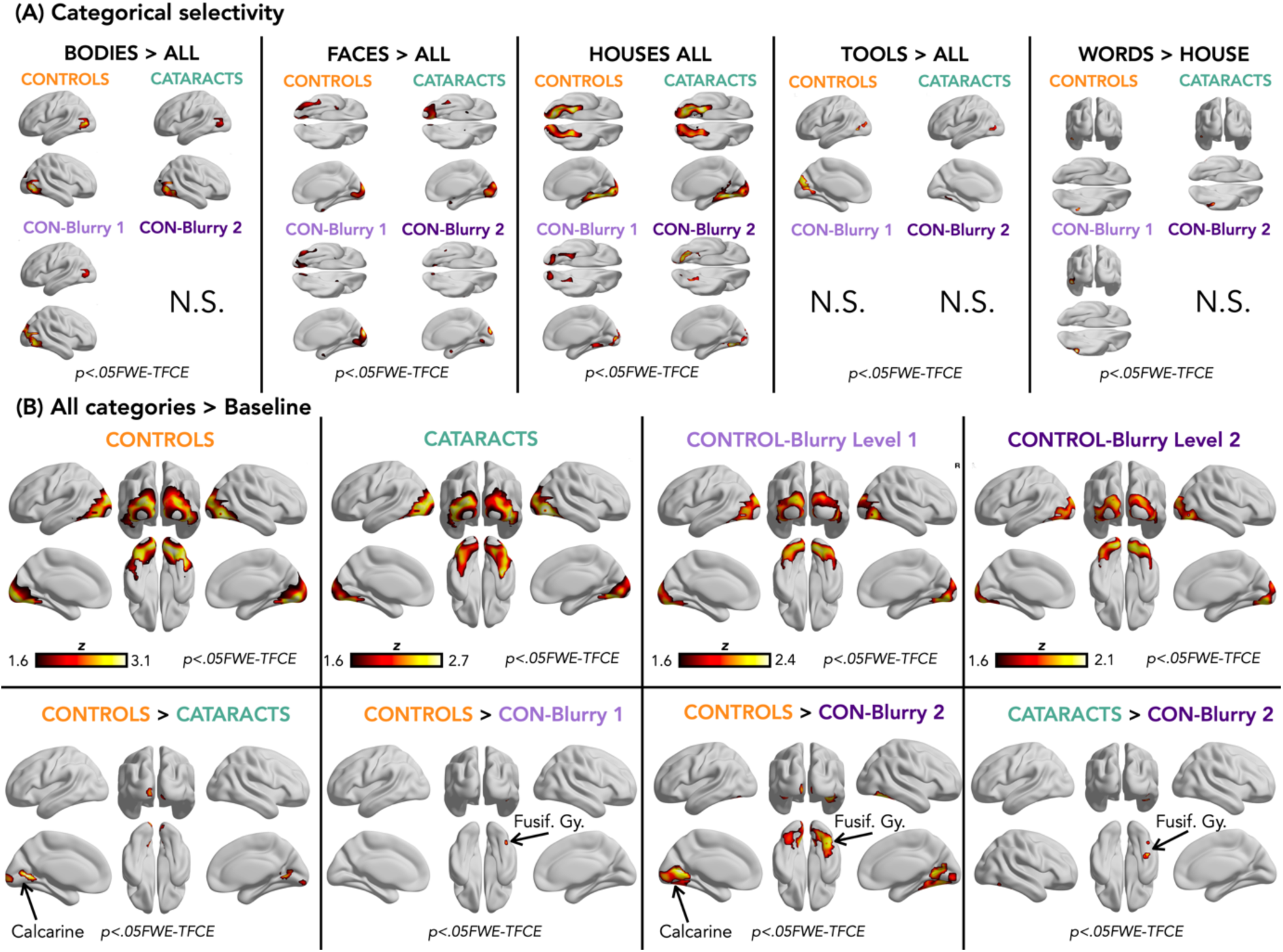
Results from Univariate Analysis. **Panel A** shows results from Univariate Analysis across category. Each column displays univariate analysis results for one category; from left to right: bodies, faces, houses, tools, and words. Univariate results for each category within each group and condition are reported, with brain maps thresholded at p < 0.05 FEW-TFCE correctred. For each category, the analysis contrasts that category against all others, except for words, where we report the contrast words > houses, as words > all did not yield significant results across groups. We did not observe any significant difference in the VOTC categorica selectivity between Controls and Cataract reversals. For more detail about the contrasts with ConBlurrry 1 and ConBlurrry 2 groups, see SI figure1. **Panel B (All stimuli > Baseline)**. The top row presents results from the univariate analysis within each group and condition: Controls, Cataract-reversals, Control-blurry level 1, and Controls-blurry level 2. The second row shows contrasts between groups that yielded to significant results. Control > Cataract, Control > Con-Blurry1, Control > Con-Blurry2 and Cataract-reversals > Con-Blurry 2. All these maps are reported with a p < 0.05 FWE and Threshold-Free Cluster Enhancement (TFCE) corrected threshold. The groups’ contrasts that are not depicted, did not show any significant difference between groups in occipito-temporal brain regions. BrainNet Viewer was used for the visualization of brain maps (Xia et al., 2013).

#### Split-half analysis

We ran a split-half analysis to investigate how reliable/stable were the patterns of activity produced by the visual stimuli in both cataract and control subjects and in the two versions of the control experiment (fig. 4).

**Figure 4:**
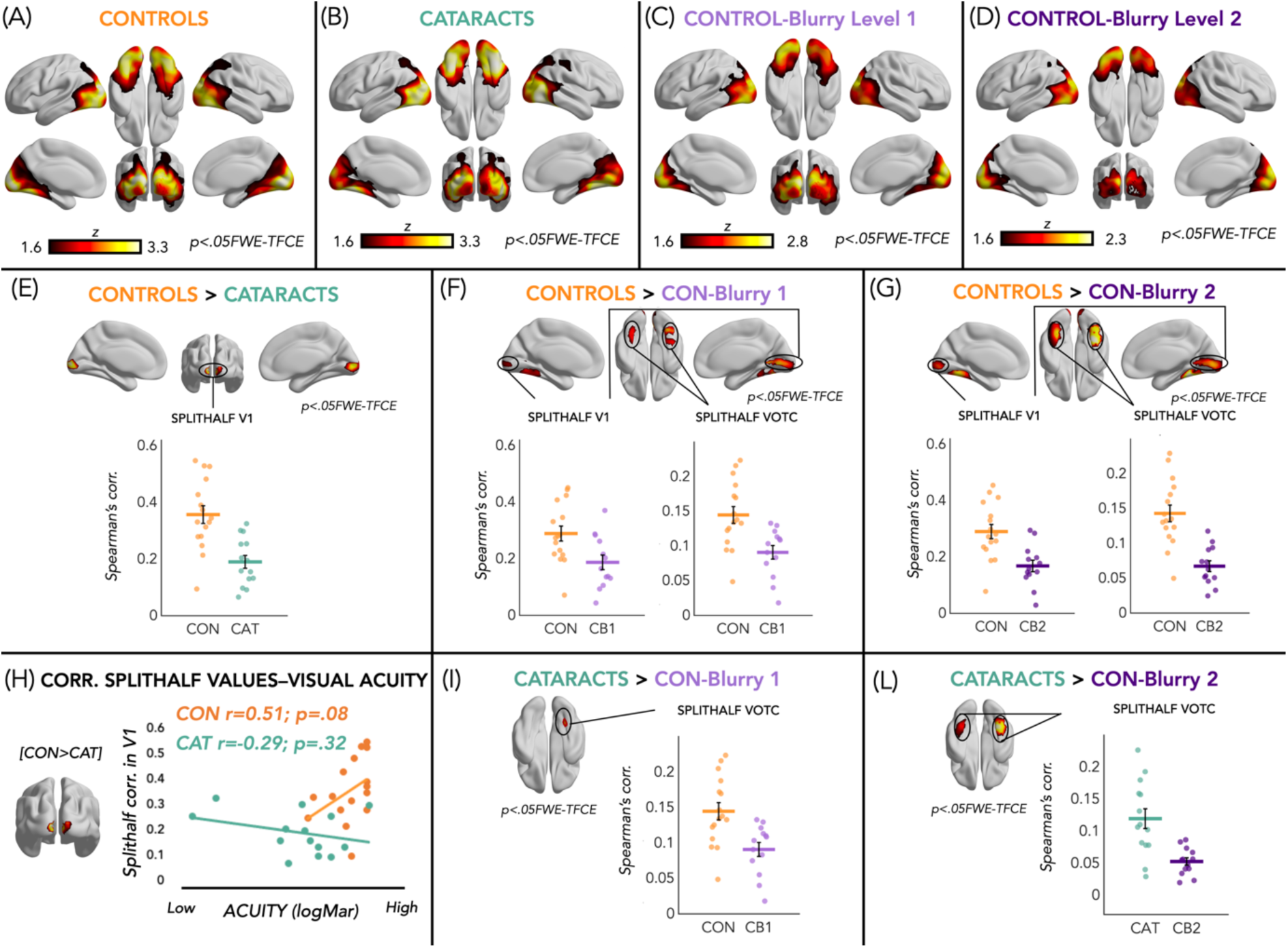
Results from Split-Half Analysis. The top row presents results from the searchlight analysis within each group and condition: controls (**Panel A**), cataract-reversals (**Panel B**), Control-blurry exp 1 (**Panel C**), and Controls-blurry exp 2 (**Panel D**). The second row shows contrasts between the Control group and the other three groups: Control > Cataract in **Panel E**, Control > Con-Blurry1 in **Panel F**, and Control > Con-Blurry2 in **Panel G. Panel H** displays the correlation between split-half values extracted from V1, where Controls show higher split-half correlation than Cataract, and visual acuity levels for each group. In **Panels I and L**, results are shown for contrasts between the Cataract-reversal group and the two control condition groups, respectively: Cataract > Con-Blurry1 in Panel I and Cataract > Con-Blurry2 in Panel L. All these maps are reported with a p < 0.05 FWE and Threshold-Free Cluster Enhancement (TFCE) corrected. For regions showing group differences, average split-half correlation values are extracted and presented as dot plots below the brain maps for visualization purposes only, without statistical analysis (as these analyses would be circular, see Kriegeskorte et a., 2008). BrainNet Viewer was used for the visualization of brain maps (Xia et al., 2013).

We split the data in two halves (i.e. odd and even runs), and we computed in each searchlight sphere (100 voxels) of the brain a value of stability of the pattern of activity produced by the images. To do so, we created for each searchlight sphere a matrix including for each stimulus the correlation between the pattern of activity from one half of the data with the pattern of activity produced by the other half of the data. In our case it is a 30*30 matrix since we have 30 images in total. Note that the on-diagonal values represent the correlation between the same images in the two different halves. Then, we computed the average of the on-diagonal values minus the average of the off-diagonal values. A positive value would indicate more similar patterns for the same stimuli (across the two halves) than across stimuli; therefore it can be considered as a measure of the stability of the brain representation elicited by discrete visual images.

For the groups’ contrasts of the split-half correlation maps (two-sample t-tests) the TFCE correction was applied within the general visual mask including both V1 and VOTC (for details about the masks, see the section above: *Statistical procedure and brain masks for TFCE correction)*, since for this analysis the difference between groups could emerge for both visual low-level and/or categorical representations.

#### Representational similarity analysis (RSA): high-level vs low level representational models

We further assessed whether the representational content encoded in the occipito-temporal cortex differed across groups using RSA (Kriegeskorte, 2008; Kriegeskorte et al., 2006). This analysis was performed using the CoSMoMVPA (Oosterhof et al., 2016) toolbox, implemented in Matlab R2016b (Mathworks). RSA is based on the concept of representational dissimilarity matrices (RDMs): a square matrix where the columns and rows correspond to the number of the conditions (i.e., n 30 image stimuli, therefore matrix of 30X30 in this experiment) and it is symmetrical about a diagonal of zeros. Each cell contains the dissimilarity index between two stimuli (Kriegeskorte & Kievit, 2013). This abstraction from the activity patterns themselves represents the main strength of RSA, allowing a direct comparison of the information carried by the representations in different groups and between brain and models (Kriegeskorte, 2008; Kriegeskorte & Mur, 2012). Crucially for the present study, RSA allowed us to use the same set of visual stimuli to isolate both the categorical/high-level and the visual /low-level representations all along the ventral occipito-temporal cortex of our participants. For each 100-voxels searchlight sphere and in each subject, we extracted the RDM, computing the dissimilarity between the spatial patterns of activity for each pair of conditions/images. To do so, we extracted the stimulus-specific BOLD estimates from the contrast images (i.e. SPM T-maps) for all the 30 image stimuli separately. Then, we used Pearson’s correlation to compute the distance (i.e.,1-r) between each pair of patterns.

Since the RDMs are symmetrical matrices, for all the RSA analyses we only used the upper triangular RDM, excluding the diagonal to avoid inflating correlation values (Ritchie et al., 2017).

Then, we used Spearman’ s partial correlation to compare the selective brain representation of each searchlight sphere with two representational models: a categorical and a low-level visual model.

The categorical RDM (fig. 5H right side) assumes that image stimuli from the same category gather together into 5 distinct clusters representing the 5 main categories (i.e. (1) bodies, (2) faces, (3) houses, (4) tools, (5) words).

**Figure 5.**
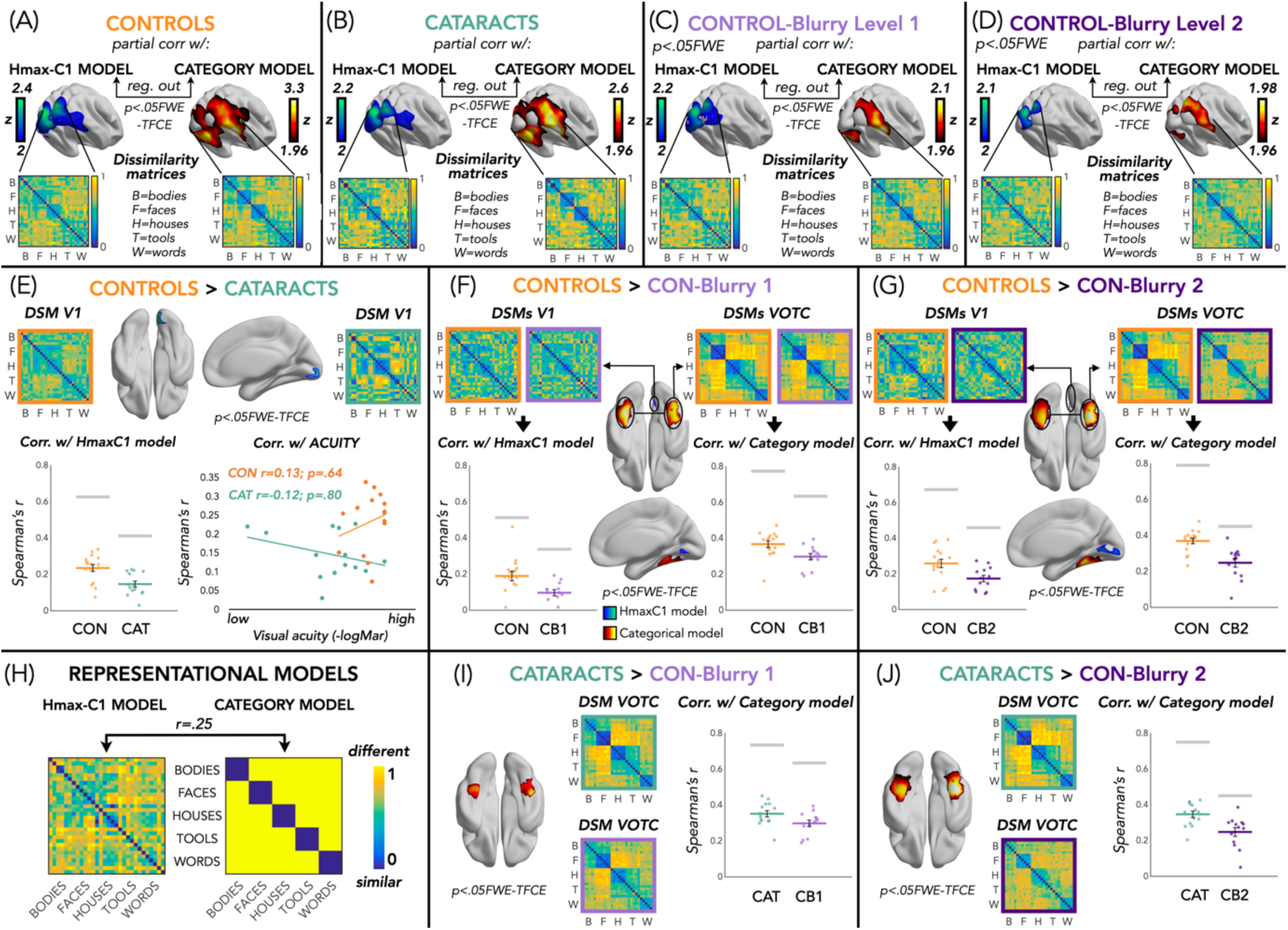
Results from Searchlight RSA Analysis with Hmax-C1 and Categorical Model. The top row presents results from the searchlight analysis within each group and condition: controls (**Panel A**), cataract-reversals (**Panel B**), Control-blurry exp 1 (**Panel C**), and Controls-blurry exp 2 (**Panel D**). Within each panel, Spearman’s r of brain dissimiarity with the Hmax-C1 model is depicted on the left, and Spearman’s r with the categorical model is depicted on the right. Below the brain maps, dissimilarity matrices extracted from the portion of the brain showing significant correlation with either the Hmax-C1 model or the categorical model are depicted. The second row shows contrasts between the Control group and the other three groups: Control > Cataract in **Panel E**, Control > Con-Blurry1 in **Panel F**, and Control > Con-Blurry2 in **Panel G. Panel H** displays the two representational models: Hmax-C1 on the left to test for low-level visual properties and the categorical model on the right to test for high-level visual representation. In **Panels I and L**, results are shown for contrasts between the Cataract-reversal group and the two control condition groups, respectively: Cataract > Con-Blurry1 in Panel I and Cataract > Con-Blurry2 in Panel L. All these maps are reported with a p < 0.05 FWE and Threshold-Free Cluster Enhancement (TFCE) corrected threshold. For regions showing group differences, brain representational dissimilarity matrices are represented from both groups, and their correlation with the HmaxC1 and the categorical models are reported as dot plots for visualization purposes only. BrainNet Viewer was used for the visualization of brain maps (Xia et al., 2013).

The Hmax model is a computational model of object recognition in the cortex that has been designed to reflect the hierarchical organization of the visual cortex (Hubel & Wiesel, 1962) in a series of layers from V1 to infero-temporal (IT) cortex (Riesenhuber & Poggio, 1999; Serre et al., 2007). To build our low-level visual model we used the output from the V1-complex cells layer (also called C1 layer; (Serre et al., 2007). The inputs for the model are the gray-value luminance images presented in the experiment. Each image is first analyzed (i.e. filtered) by an array of simple cells (S1) units at 4 different orientations and 16 scales. At the next C1 layer, the image is subsampled through a local Max pooling operation over a neighborhood of S1 units in both space and scale, but with the same preferred orientation (Serre et al., 2007). C1 layer stage corresponds to V1 cortical complex cells, which show some tolerance to spatial shift and size (Serre et al., 2007). The outputs of all complex cells were concatenated into a vector as the V1 representational pattern of each image (Khaligh-Razavi & Kriegeskorte, 2014; Kriegeskorte, 2008). We, finally, built the (30 × 30) RDM computing 1-Pearson’s correlation of each pair of vectors (fig. 5H left side). The Hmax-C1 RDM was significantly correlated with the categorical RDM (r = 0.25, p<0.001). The use of partial correlation between the brain RDMs and the models, allowed us to regress this shared correlation between the two models and to look at the unique correlation between the brain RDM and each model independently.

For each model, the output correlation values with the brain RDM were Fisher transformed and assigned to the center voxel of each searchlight sphere. We, therefore, obtained two separate correlation brain maps, one for each model. To estimate a group-level statistic we performed a voxel-wise t-test against baseline, for each group separately: the control group, the cataract-reversal group, the CON-blurry1 group and the CON-blurry2 group.

We also performed two-sample t-tests to check for differences between groups. For these groups’ contrasts the TFCE correction was applied within the V1 mask for the correlations with the HmaxC1-model and within the VOTC mask for the correlations with the categorical model (for details about the masks, see the section above: *Statistical procedure and brain masks for TFCE correction)*. This choice was guided by our a priori expectation— also supported by our within-group whole-brain results (fig. 5 panels A-B-C-D) —regarding where these two models would be most represented in the brain (i.e., HmaxC1 in V1 and the categorical model in VOTC).

#### Decoding analysis between and within categories

We complemented the RSA analyses with two different decoding analyses, one targeting the categorical representation in VOTC (5-way categories decoding) and one targeting the low-level visual representation in V1 (6-way items decoding for each category). These analyses were performed using the CoSMoMVPA (Oosterhof et al., 2016) toolbox, implemented in Matlab R2016b (Mathworks).

The 5-way categories decoding analysis is similar, on the theoretical level, to the RSA analysis testing the relation between brain representation and the categorical model. In both analyses, we expect that in VOTC, items from the same category have more similar patterns of activity compared to items from different categories. To foreshadow our results, since we observed no differences across control and cataract-reversals groups in the relation between brain representation in VOTC and the categorical model, we decided to carry out the between-category decoding analysis to corroborate our observation of no differences across these groups in how VOTC encodes categories of visual stimuli. However, given the conceptual redundancy of these analyses we placed the 5-way decoding between categories in the supplemental material (SI fig. 2).

The 6-way item decoding within (fig. 6) each category is an analysis that will, instead, add complementary information to the RSA analysis with the HmaxC1 model.

**Figure 6.**
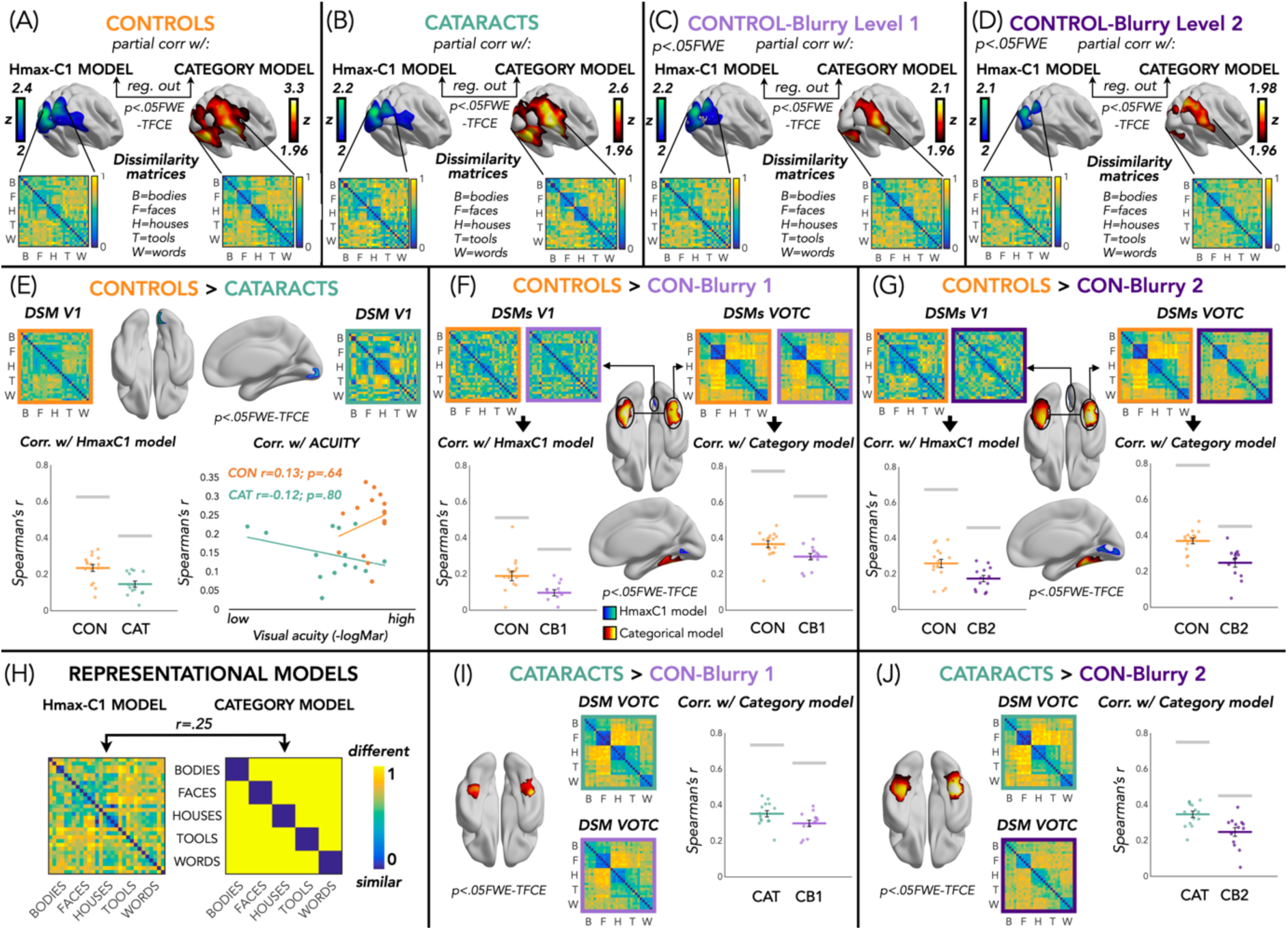
Within-category decoding analyses. These results comprise five different decoding analyses, each executed on a subset of the data, focusing on one category at a time. Each column in the figure represents the analysis for a specific category: bodies in **panel A**, faces in **panel B**, houses in **panel C**, tools in **panel D**, and words in **panel E**. The top row (**panels A1, B1, C1, D1 & E1**) presents, for each category, the six images belonging to that category alongside brain maps illustrating the regions of the brain exhibiting significant decoding accuracy for classifying those six images within each group. The second row (**panels A2, B2, C2, D2, E2**) displays contrasts between the Control group and the Cataract-Reversal group for each category individually. The third row (**panels A3, B3, C3, D3, E3**) exhibits contrasts between the Control group and the Control-Blurry 1 condition for each category separately, while the fourth row (**panels A4, B4, C4, D4, E4**) demonstrates contrasts between the Control group and the Control-Blurry 2 condition for each category individually. All brain maps are presented with a threshold corrected for Family-Wise Error (FWE) and Threshold-Free Cluster Enhancement (TFCE) at p < 0.05. Decoding accuracies for regions showing group differences are visualized using dot plots for visualization purposes only. BrainNet Viewer was used for the visualization of brain maps (Xia et al., 2013).

In our study the selection of the categorical model was based on our own design (we created a stimulus set with 5 well defined categories). For the low-level visual model, we selected the HmaxC1 model that is well known to fit well the representation implemented in V1 (Riesenhuber & Poggio, 1999; Serre et al., 2007). However, there are many visual features that we could have tested separately (e.g. rectilinearity/curvilinearity; simple silhouette model; gist model; etc.), with the downside of inflating the multiple comparisons problem. But the 6-way item decoding does not require an a-priori selection of the low-level feature or model to be tested. In this case, in fact, the decoding algorithm will select by itself the features to use to classify the items. In addition, since this analysis is implemented within each category (we run 5 independent analyses within 5 different subsets of data, one for each category), we circumvent the confound of partial collinearity between categorical membership and low-level features (e.g. all faces share similar visual properties).

In all the classification analyses (between- and within-categories) we ran multiclass decoding analyses. In the decoding analysis between categories (5-way category decoding) we tested the discriminability of patterns for the five categories using a linear discriminant analysis (LDA). Since categories are known to be represented in VOTC, we expected that this task would be mostly performed in VOTC. In the decoding analyses within each category (6-way items decoding for each category) we tested the discriminability of brain voxel activation patterns for the six items within each category, using LDA. We run 5 separate analyses, one for each category. Since the low-level visual features of the items within each category were quite variable, we expected that V1 would mostly engage in such image specific decoding (e.g. the classification of the 6 images of bodies would be mostly based on the different low level visual properties of the 6 bodies) (Badwal et al., 2024). We, therefore, obtained five separate accuracy maps, one for each category.

In all these decoding analyses, we performed a leave-one-run-out cross-validation procedure using beta-estimates from 4 runs in the training set, and the beta-estimates from the remaining independent run to test the classifier, with iterations across all possible training and test sets. This procedure was implemented in each 100-voxels sphere using a searchlight approach (Kriegeskorte et al., 2006; Tong & Pratte, 2012). Classification accuracy for each sphere was assigned to the central voxel of the sphere to produce accuracy maps. The resulting accuracy maps were then smoothed with an 8-mm Gaussian kernel.

To estimate a group-level statistic we performed a voxel-wise t-test against chance level, for each group separately: the control group, the cataract-reversal group, the CON-blurry1 group and the CON-blurry2 group. We also performed two-sample t-tests to check for differences between groups. For the groups’ contrasts in the between-categories decoding analysis, the TFCE correction was applied within the VOTC mask since we expect this kind of classification to be based on categorical membership. For the groups’ contrasts in the within-category decoding analyses instead, the TFCE correction was applied within the V1 mask because, as mentioned above, we expected that V1 would mostly engage in such image specific decoding. For details about the masks, see the section above: *Statistical procedure and brain masks for TFCE correction*.

#### Deep Neural Network Analyses

Earlier studies comparing representations between Deep Neural Networks (DNNs) and the human brain revealed a spatio-temporal hierarchical alignment: as DNN layer number increased, DNN representations exhibited stronger correlation with cortical representations emerging later in time, and in progressively downstream brain regions along both the dorsal and ventral visual pathways (Cichy et al., 2016). We leveraged this alignment to gain further insights into the mechanisms underlying visual information processing after transient blindness early in life (Celeghin et al., 2023).

##### Model Architecture

To explore the alignment between deep neural network (DNN) outcomes and human functional magnetic resonance imaging (fMRI) data, we used AlexNet (Krizhevsky et al., 2017). The choice of AlexNet was driven by its significance in image recognition tasks and its architectural design, which is well-suited for representing the hierarchical visual processing observed in the human visual cortex. Despite clear differences between humans and DNNs (Geirhos et al., 2018; Kell et al., 2018; Kriegeskorte, 2015; Maniquet et al., 2024), prior research has indicated that the early layers of convolutional neural networks, like those in AlexNet, develop features tunings (e.g., oriented spatial frequency detectors) and representational geometry that are similar to those observed in the primary visual cortex (Cadieu et al., 2014; Guclu & Van Gerven, 2015; Khaligh-Razavi & Kriegeskorte, 2014), while the deeper layers have shown representational alignment with high-level ventral visual areas such as IT (Khaligh-Razavi & Kriegeskorte, 2014) which are known to process more complex categorical information. These characteristics make AlexNet a good model for investigating the potential parallels between artificial and biological visual systems in low- and high-level areas.

##### Data Preparation

To train our models, we utilized the ImageNet dataset (Russakovsky et al., 2015) which includes 1000 diverse image categories. This dataset is provided in two forms: a standard version and a modified "blurred" version to simulate the effects of visual impairments such as cataracts.

In the blurred version of the dataset, a low-pass Butterworth filter was applied to each image. This was implemented using the ‘filter.butterworth’ class from the scikit-image 0.23.2 library (Van Der Walt et al., 2014) in Python 3.10. We conducted tests with varying levels of blur to identify the specific level that degrades the representation in the V1-like layer (the lower layer of the DNN) in a manner consistent with fMRI findings in cataract-reversal and control-blurry groups in the V1 brain area. The final filter, configured with an order of 2 and a cutoff frequency ratio of 0.04, attenuates high-frequency components. This modification mimics the blurring effect observed in cataract-impaired vision, generating a realistic dataset for assessing our model robustness to such visual impairments. Note that it does not mimic the vision before cataract removal, when vision can be so poor that acuity is even not measurable.

During the training and validation phases, both the standard and blurred ImageNet images underwent several transformations to enhance model robustness and prevent overfitting. For training, images were randomly resized and cropped to dimensions of 224×224 pixels and subjected to random horizontal flips to ensure variability. These images were then normalized using specified mean values of [0.485, 0.456, 0.406] and standard deviations of [0.229, 0.224, 0.225]. Similarly, validation images were resized to a uniform shorter dimension of 256 pixels before being cropped to 224×224, and underwent the same normalization process to maintain consistency in image processing across different datasets.

In the testing phase, the same 5-categories dataset used for human participants (see stimuli description in the Methods section) was employed to extract the networks’ activations, again in both standard and blurred versions to simulate, respectively, normal and cataract visual properties. For this phase, the images were resized to 224×224 pixels and normalized using the same mean values and standard deviations as the training and validation images.

##### Training and Testing Conditions

Three distinct training and testing paradigms were designed to model the different visual processing conditions experienced by our human participants (see the section about participants and control experiment for a detailed description of these conditions in the fMRI experiment):

Intact-to-Intact (‘Control’ Network): In this baseline condition, the network was both trained and tested on standard, unaltered images. This setup served as the control to assess the network’s performance under normal visual conditions.

*Blurry-to-Blurry* (‘Cataract’ Network): For this condition, both training and testing phases involved images that had been previously blurred to simulate cataract-like visual impairments. The blurring was applied as detailed in the data preparation subsection. This approach was designed to mimic the visual degradation typical in cataract-affected vision, allowing us to study the impact of such impairments on the network’s performance and internal representations.

Note that the blurring only mimics vision after cataract removal, and not the near-blindness levels prior to the surgery. Based on the literature described in the introduction, we know that this pre-surgery deprivation will be particularly detrimental for the development of responses in early visual cortex due to the existence of an early sensitive period. To mimic this aspect of the development in the cataract patients in at least a crude manner, the weights of the first convolutional layer were frozen (i.e., not trained) during the training phase of the Cataract-like network.

*Intact-to-Blurry* (‘Control-Blurry’ Network): Here, the network was trained on standard, unaltered images but tested on blurred ones. This paradigm simulates the experience of the control-blurry conditions, where individuals with normal vision were presented with blurred images. The application of the Butterworth filter for testing mirrors the procedure used in the Blurry-to-Blurry condition.

##### Features Extraction

All the DNN activations in this study were extracted from the layers ‘features.1’ and ‘classifier.5’ during the testing phase on our 5-categories dataset. Specifically, features.1 is the activation (ReLU) layer after the first convolutional layer, chosen to capture early, low-level representations of the data. Henceforth, features.1 will be referred to as the V1-like layer. In contrast, classifier.5, the activation layer following the second-last linear layer, was selected to capture late, high-level representations, analogous to the higher visual and categorical processing observed in VOTC. This layer will be referred to as the VOTC-like layer. The selection of these layers allows for a direct comparison between the neural network’s processing stages and distinct functional areas of the brain, thus providing insight into how artificial systems can mimic biological visual processing.

To align our DNN analysis with the multivariate approaches used on brain data, which included 5 runs per subject, we introduced artificial variations into the DNN activations. Specifically, we generated 5 "runs" or variations by applying Gaussian noise with a mean of 0.0 and a standard deviation of 6 to the networks’ activations. This level of noise was chosen to result in a comparable overall classification performance in the control network relative to the human control group.

#### Direct correlation of RDMs from the brain ROIs with RDMs extracted from DNN layers (in the intact-to-intact DNN condition)

This initial analysis pursued a dual objective. Firstly, it aimed to demonstrate that the DNN, trained and tested with typical stimuli, along with the two specific layers we identified, indeed depicts a representation of our stimuli consistent with our biological data. Secondly, it sought to enhance and expand our RSA results by employing representational models derived from RDMs extracted from the DNN network, thereby minimizing a priori bias in model selection.

In this analysis, we generated RDMs from the activations of the V1-like and VOTC-like layers of the Control network using a Pearson’s correlation distance metric. This network was specifically trained and tested on normal (non-blurred) images, serving as an analogue to our human control group. We then calculated Spearman’s correlations between the RDMs from these network layers and the brain-derived RDMs obtained from the V1 and VOTC ROIs across all participant groups and conditions (Controls, Cataract-reversals, Con-blurry1, and Con-blurry2).

To run statistics on these data we performed the following non-parametric tests: Wilcoxon test for one sample t-tests to check significance against zero and for paired sample t-tests to check differences of brain/layers correlation within the same groups; Mann-Whitney tests for the statistical comparisons between groups. To address multiple comparisons, all p-values were corrected using false discovery rate (FDR) (Benjamini & Hochberg, 1995).

#### RSA analyses: correlation of RDMs extracted from different layers and conditions of DNN with representational models (HmaxC1 and categorical)

After demonstrating that the two layers in the DNN, trained and tested with typical images, accurately simulated the processing hierarchy we observe in our brain data, we proceeded to conduct the same RSA analysis previously performed on the brain data. This involved using RDMs extracted from the three different DNN conditions. In this analysis, RDMs were extracted from the V1-like and the VOTC-like layers across the various training and testing conditions mimicking the visual conditions in our human control, cataract and control-blurry groups.

Based on the results from our biological data, we conducted the following analyses: for each set of RDMs extracted from the various DNN conditions from the V1-like layer, we computed Spearman’s correlation with the HmaxC1 model; whereas for the RDMs extracted from the different DNN conditions from the VOTC-like layer, we computed Spearman’s correlation with the categorical model. We controlled for the partial correlation between the two models.

We implemented non-parametric permutations to conduct statistical analyses. For each layer/condition, we determined the statistical difference from zero using a permutation test with 10,000 iterations. This involved constructing a null distribution for the correlations by computing them after randomly shuffling the conditions’ labels.

Additionally, we assessed the statistical difference between each group-like condition using a permutation test. We generated a null distribution for the difference in correlation values of the two conditions by computing the difference after randomly shuffling the conditions’ labels, repeating this step 10,000 times.

To determine the statistical significance of our results, we compared the observed result to the null distribution. This comparison involved calculating the proportion of observations in the null distribution that had a value higher than the one obtained in the real test. To address multiple comparisons, all p-values were corrected using false discovery rate (FDR) (Benjamini & Hochberg, 1995).

#### Between- and within-category decoding analyses in different layers and conditions of the DNN

For each DNN condition, we replicated the decoding analyses previously conducted on the brain data. We followed the exact procedure outlined for the fMRI data (see ‘decoding analysis’ in the methods section) with the sole distinction being that, rather than using brain activity patterns, we employed activations extracted from both the V1-like and the VOTC-like layers. Furthermore, we restricted the decoding analyses to the analyses of interest based on results from our biological data: we conducted between-category decoding in the VOTC-like layer and within-category decoding in the V1-like layers. To conduct statistical analyses, we implemented the same non-parametric permutations procedure described in the previous section.

## Results

### Visual acuity

The control group has a mean level of logMar of 0.01 (SD 0.07, min: -0.05, max:0.17), the cataract-reversal group (see table 1) has a mean level of logMar of 0.21 (SD 0.18, min - 0.05, max: 0.61). As expected (Lewis et al., 1995; Sinha & Held, 2012; Zohary et al., 2022), an independent samples T-test revealed a significant difference (fig. 2A) of the level of visual acuity between the two groups (t(28)=–4.1; p=0.0003).

### Behavioral response in the fMRI

During the fMRI data acquisition, participants were asked to identify instances where two images did not repeat. We excluded one subject in the control group, one subject in the cataract-reversal group and one subject in the control experiment (CONblurry1 and CONblurry2) based on poor performance during fMRI data acquisition (under 2.5 standard deviations from the group mean).

After the exclusions, the mean values of accuracy were 94% in the control group, 91% in the cataract-reversal group, 94% in the CON-blurry1 experiment, and 95% in the CON-blurry2 experiment (fig. 2B). Independent samples t-tests did not show any significant difference between the accuracy values across groups (pFDR between 0.63 and 0.99).

### Amount of gaze displacement

Ten out of the 15 cataract-reversal subjects reported experiencing nystagmus (Table 1), a comorbid condition often associated with early visual deprivation and characterized by involuntary and repetitive eye movements. The small tracking window of typical in-scanner eye trackers in combination with the fact that participants wore individually fitted in-scanner corrective glasses (the same glasses that they wore during the visual acuity test) and the presence of nystagmus made the use of eye-tracker impracticable. We instead employed deepMReye (Frey et al., 2021) to examine the eye movement patterns of our subjects and used this data to account for this potential confounding factor in our study (fig. 2C).

The average amount of gaze displacement inside the scanner, computed through deeepMReye, was significantly higher in the cataract-reversal group compared to all the other groups (fig. 2C; CAT vs CON (t(28)=2.61; p=0.01; pFDR=0.02); CAT vs CON-blurry1 (t(25)=-3.1; p=0.004, pFDR=0.02); CAT vs CON-blurry2 (t(25)=-2.78; p=0.01, pFDR=0.02). Instead, the amount of gaze displacement did not differ between the control group and the CON-blurry1 (t(27)=0.15; p=0.88, pFDR=0.94), nor between the control group and the CON-blurry2 (t(27)=0.08; p=0.94, pFDR=0.94).

This difference between the caract-reversal group and the others was expected, due the presence of nystagmus in most of the cataract-reversal individuals, and validates the indices provided by deepMReye. As described in the method section, we included the variance for the X and Y gaze positions as regressors of no-interest in our GLM (see next paragraph), to control for the impact of different magnitude of eye movement on the fMRI data activity.

### Univariate analysis

We conducted this analysis to examine the activated regions (within and between groups) for all stimuli compared to the baseline and for each category in contrast to all other categories. It is crucial to note that our experimental design was optimized for multivariate analyses, and the rapid event-related presentation typically does not yield the best signal for univariate analysis.

Overall, univariate categorical preference in VOTC was found in the expected locations (Rosenke et al., 2021) in cataract-reversal and control groups with only minor differences across groups (figure 3A). Most importantly, we did not observe lower categorical selectivity in the cataract-reversal group when compared to controls in VOTC (figure 3A and SI fig1). Even using a lenient threshold of p= .001 uncorrected for multiple comparisons did not reveal any difference between the cataract and the control groups for category selective responses, reinforcing the true absence of differences across groups (See supplemental figure 1).

Since we did not observe any difference in the preferential response to any of our 5 categories, we additionally carried out a univariate contrast between groups when collapsing all stimuli together to investigate potential difference in the global visual response to all visual stimuli (Fig. 3B). As expected, all groups (fig. 3B, first row) exhibit significant activation for visual stimuli across all categories compared to the baseline, covering a broad area of the visual cortex, including both the early visual cortex and more anterior occipito-temporal regions. However, when looking at the between groups results using independent sample T-tests, some differences emerge. The significant contrasts from these groups’ comparisons are illustrated in fig. 3B (second row). The comparison between controls and the cataract-reversal group revealed less activation in a portion of V1 in the cataract-reversal group, with no difference observed in VOTC between the two groups. Comparing controls with controls-blurry1 highlighted only a small region of the posterior fusiform gyrus with lower activity in controls-blurry1 compared to the control group. We also observed decreased activation in controls-blurry2 compared to controls, encompassing a broad area of both V1 and the fusiform gyrus. Additionally, a lower activation was observed in a portion of VOTC in controls-blurry2 compared to the cataract-reversal group.

These results, along with those concerning specific category contrasts, do not reveal any significant differences in the activity of VOTC in cataract-reversal individuals compared to controls. Additionally, the findings from comparisons with control-blurry 1 and control-blurry 2 conditions are not always straightforward and are challenging to interpret (see SI fig1). Therefore, we conducted a series of multivariate pattern analyses to further explore potential differences in categorical representation in VOTC between cataract-reversal individuals and controls, as well as to better elucidate the representation in the control-blurry 1 and control-blurry 2 conditions.

### Split-half analysis

We conducted this analysis to investigate where in the brain visual stimuli trigger a response that discriminates between the stimuli in a consistent and stable manner, and to assess whether this stability is similar across different groups. The procedure involves examining data consistency between two distinct portions (i.e., two halves) of the dataset. The brain regions that exhibit highly similar activation patterns in both halves are the ones that respond consistently and meaningfully to our stimuli. Since our participants were presented with visual stimuli, we anticipate that the visual cortex will predominantly represent the images in a reliable manner.

In fig. 4, the first row displays brain maps resulting from the whole-brain searchlight split-half analysis for each group individually (panels A, B, C, D).

As expected, all the groups (fig. 4, panels A-D) show high reliability in the representation of visual stimuli in occipital regions, encompassing both EVC and VOTC.

The comparison (independent sample t-tests 2-tailed) between controls and cataract-reversals showed lower split-half correlation in the cataract-reversal group in V1, with no between-group differences in VOTC (fig. 4E). In contrast, reduced split-half correlation in both V1 and VOTC was observed when comparing controls-blurry1 (fig. 4F) and controls-blurry2 (fig. 4G) to controls without burring. Finally, when we directly compared the cataract-reversal group with the controls-blurry1 (fig. 4I) and with the controls-blurry2 (fig. 4L) we did not observe any difference in V1 but significantly lower split-half correlation in VOTC for both controls-blurry1 and controls-blurry2 when compared with the cataract-reversal group.

### Representational similarity analysis (RSA): low vs high-levels representational models

Representational similarity analysis enabled us to simultaneously probe the low-level and categorical representations throughout the ventral occipito-temporal cortex in our participants. We correlated the brain representation of our stimuli with two representational model based either on the low-level visual features of the images (i.e. HmaxC1 model) or on their categorical membership (see fig. 5H).

The resulting maps from the whole brain searchlight RSA analyses for both models and in each group separately are represented in the first row of fig. 5 (panels A, B, C, D).

As expected, the HmaxC1 model elicits the higher correlation with the early visual cortex, while the categorical model is significantly correlated with VOTC activity in all groups (fig. 5, panels A-D).

The comparison between controls and cataract-reversals showed a significantly lower correlation in cataract-reversals with the HmaxC1 model only in V1, while there was no group differences in the correlation with the categorical model in VOTC (fig. 5E).

In both blurry1 and blurry2 control groups, there was a reduced correlation with the HmaxC1 model in a big portion of V1 compared to the control group and a lower correlation with the categorical model in VOTC (see fig. 5F-G).

Finally, when we directly compared the cataract-reversal group with the controls-blurry1 (fig. 5I) and with the controls-blurry2 (fig. 5J) we did not observe any difference in the correlation with the HmaxC1 model in V1. However, we did find a significantly lower correlation with the categorical model in VOTC for both controls-blurry1 and controls-blurry2 when compared with the cataract-reversal group.

### Decoding analyses

We expanded the RSA analyses by incorporating two additional decoding analyses. One of these analyses focused on decoding categorical representation in VOTC. This analysis was implemented to strengthen our observation of no group differences using RSA using a categorical model of stimuli encoding (see fig. 5). However, whereas RSA aims to identify maximal relations, decoding seeks to pinpoint maximal distances between patterns and, in this instance, categories. The results from the decoding analysis closely matched those obtained from the RSA analysis, providing strong support for our observation of no differences between the cataract and control group in how categories are encoded in VOTC. Due to the conceptual overlap of these findings, we decided to present the 5-way decoding between categories analysis in the supplemental material (see SI fig. 2).

The second addition is the 6-way item decoding, applied within each category, which complements the RSA analysis with the HmaxC1 model. As we conducted five separate analyses, decoding the items within each category and including only the six items from the same category at a time, we anticipate that the algorithm’s decision-making process will rely mostly on low and middle visual properties (since the categorical membership is identical between the 6 decoded items). Our observations confirm this expectation, as we consistently found that V1 is the primary brain region encoding our stimuli across the 5 categories tested (separately) and all groups (see fig. 6, top row). You can also refer to Supplementary fig. 3 for an overlay of the 5 categories, demonstrating that the results are consistently similar across the different categories. These maps resulted from the whole brain searchlight 6-way decoding analyses for each group and for each category separately. No other brain regions were found to be involved in this task, even when considering lower p values.

Interestingly when we contrast the results from different groups with independent sample T-tests, clear differences emerged. Contrasts producing significant results are reported in fig. 6 (panels A 2-4; B 2-4; C 2-4; D 2-4; E 2-4).

The comparison between controls and cataract-reversals revealed significantly lower decoding activity in V1 among cataract-reversal individuals compared to controls. This consistent pattern of results was evident across all categories: bodies (fig. 6, Panel A2), faces (fig. 6, Panel B2), houses (fig. 6, Panel C2), tools (fig. 6, Panel D2), and words (fig. 6, Panel E2). These findings align with the results obtained from the split half analysis and RSA with the HmaxC1 model, providing robust evidence that individuals who underwent cataract reversal exhibit permanent impaired representation of low-level visual properties in V1. Indeed, they demonstrate that their primary visual cortex faces challenges in utilizing these visual features to differentiate items within the same category.

Following a similar procedure to the previous analyses, we conducted comparisons between controls and the two control conditions: controls-blurry1 (fig. 6, third row) and controls-blurry2 (fig. 6, fourth row). In both control conditions (blurry1 and blurry2), V1 exhibited reduced decoding accuracy compared to the control group. These results were consistent across all categories and closely resembled the findings observed in the contrast between controls and cataract-reversals. Direct comparisons between the cataract-reversal group and the control-blurry1 & control-blurry2 groups did not reveal any significant differences.

### Correlation analyses between Visual Acuity and brain data

To explore whether the level of visual acuity at the time of testing could account for the group differences that we observed in the fMRI analyses, we conducted correlation analyses between visual acuity values (i.e -logMar) and V1. These correlations were performed for all the analyses that showed differences between controls and cataract-reversals, including the split half analysis (fig. 4H), RSA with the HmaxC1 model (fig. 5E), and the five within-category decoding analyses. Below we report the p values FDR corrected for 7 comparisons (i.e. the total number of correlation analyses we ran). Overall, the results indicate that there were no significant correlations between brain regions showing altered response in the cataract-reversal people and visual acuity. For each of these correlation analyses, we computed the correlation between visual acuity and the results extracted from the V1 portion showing a difference in each specific analysis.

For the split-half analysis, the Spearman’s correlation between visual acuity and V1 correlation for the control group was r=0.51 (pFDR=0.084), and for the cataract-reversal group, it was r=-0.29 (pFDR=0.32), (see fig. 4H).

Similarly, for the RSA with the HmaxC1 model, the Spearman’s correlations were r=0.13 (pFDR=0.64) for the control group and r=-0.12 (pFDR=0.8) for the cataract-reversal group (see fig. 5E).

Regarding the within-decoding analyses, the Spearman’s correlations between visual acuity and V1 decoding accuracy were as follows: for bodies r=0.57 (pFDR=0.065) for the control group and r=0.21 (pFDR=0.8) for the cataract-reversal group; for faces r=0.5 (pFDR=0.065) for the control group and r=-0.04 (pFDR=0.8) for the cataract-reversal group; for houses r=0.42 (pFDR=0.13) for the control group and r=-0.15 (pFDR=0.8) for the cataract-reversal group; for tools r=0.67 (pFDR=0.04) for the control group and r=-0.25 (pFDR=0.8) for the cataract-reversal group; for words r=0.51 (pFDR=0.065) for the control group and r=-0.1 (pFDR=0.8) for the cataract-reversal group.

Overall, the within-group correlations do not suggest that the acuity at the time of test can explain the between-group differences in fMRI data, in particular because there is a total lack of such correlation in the cataract group that has the highest variance in acuity.

### DNN – AlexNet results

#### Direct correlation of RDMs from the brain ROIs with RDMs extracted from DNN layers (in the intact-to-intact DNN condition)

Earlier studies comparing Deep Neural Networks (DNNs) and human brain representations found a spatio-temporal alignment: as DNN layer depth increased, correlations strengthened with later-emerging cortical representations in downstream regions along both dorsal and ventral visual pathways (Cichy et al., 2016). Leveraging this alignment, we explored visual processing mechanisms after early-life transient blindness (Celeghin et al., 2023). Specifically, we asked whether a neural network modeled after cataract-like condition would show impairment in low-level (V1-like) visual representations while preserving high-level (VOTC-like) categorical representations—and whether a network mimicking control-blurry condition would exhibit impairments across both layers.

Results from this analysis are depicted in fig. 7B. Overall, the results show that the DNN trained and tested with typical stimuli accurately model our brain data.

**Figure 7.**
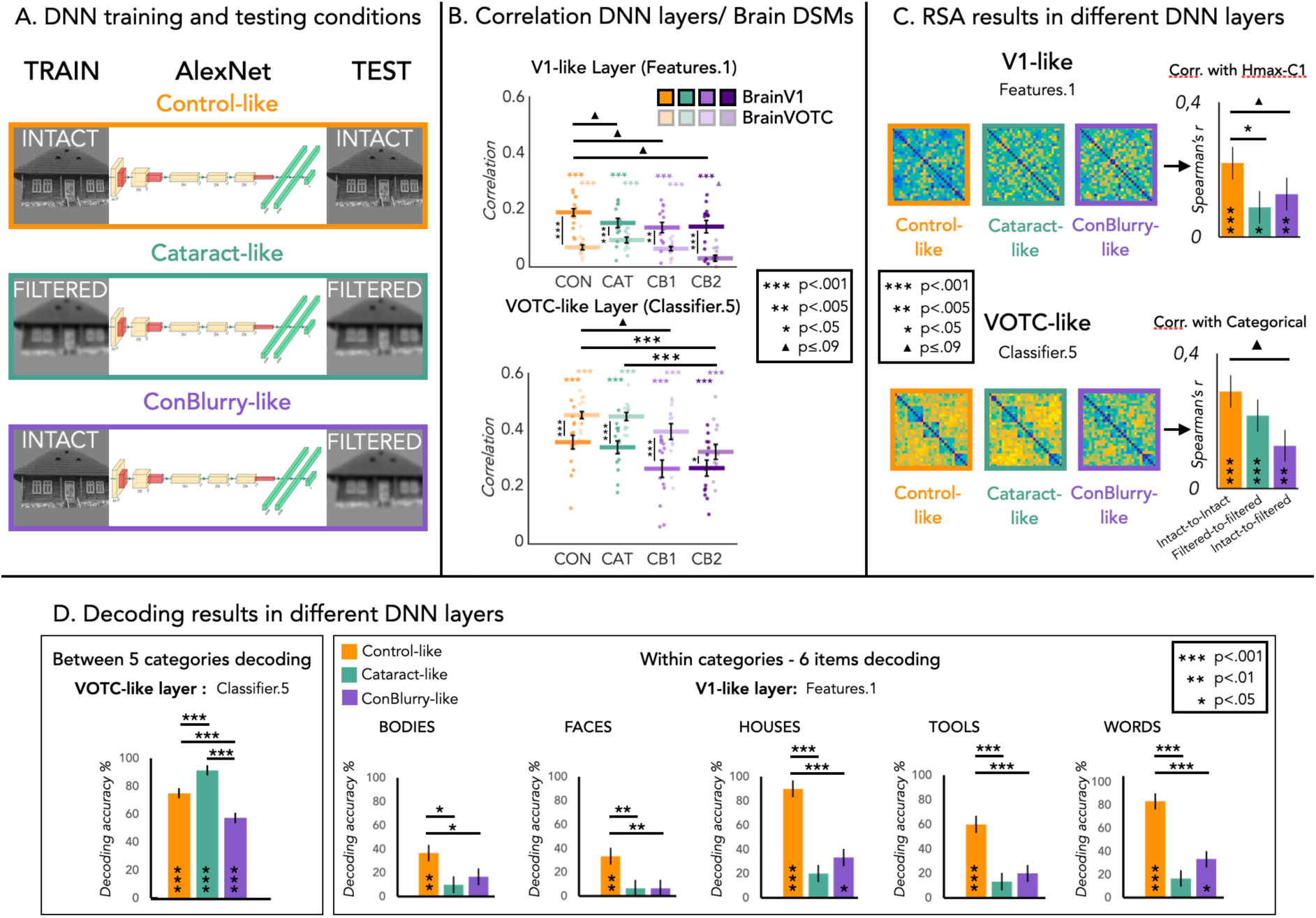
DNN – AlexNet analysis and results. In **panel A** the training and testing conditions implemented in the AlexNet are depicted. In the first row, both the training and testing phases were conducted with intact images, mimicking the control condition situation. In the second row, the training and testing phases were both conducted using filtered images to simulate the cataract-reversal condition. Finally, in the third row, the training was done using intact images, but in the testing phase, filtered images were presented, mimicking the scenario of controls who participated in the fMRI study with degraded images. In **panel B**, the correlations between the RDMs extracted from the DNN layers (trained and tested with typical images) and the RDMs extracted from brain regions in the four groups/conditions are plotted. The top section illustrates the correlation values between the RDMs extracted from the V1-like layer and those extracted from V1and VOTC in Controls, Cataract-reversals, Controls Blurry 1, and Controls Blurry 2. The bottom section displays the correlation values between the RDMs extracted from the VOTC-like layer and those extracted from V1 and VOTC in the same groups/conditions. The RSA results in V1-like and VOTC-like layers are reported in **panel C**. For both layers and for each group we depict the representational dissimilarity matrices (RDMs) and we report in the bar plots the correlation values between V1-like layer’s RDM and the HmaxC1 model and between VOTC-like layer’s RDM and the categorical model. In **panel D**, decoding results for the V1-like and VOTC-like layers are reported. In the box on the left side, the bar plots represent the results for the between categories decoding in the VOTC-like layer. In the right box, the results from the 5 within category decoding in the V1-like layers are depicted. Asterisks indicate significant values computed with permutation tests (10,000 randomizations of stimulus labels), and error bars indicate SD computed by bootstrap resampling of the stimuli.

Specifically, the RDM extracted from the first layer (V1-like layer) of the DNN exhibited a significant correlation with the RDMs extracted from the V1 ROI in each group/condition (Spearman’s r: Controls 0.171, Cataract-reversals 0.147, Controls-Blurry1 0.131, and Controls-Blurry2 0.134; all p < 0.001 at the Wilcoxon one-sample test, FDR corrected). Additionally, as expected, within each group, this correlation value was significantly higher compared to the correlation between the RDM extracted from the same first layer (V1-like layer) of the DNN and the RDM extracted from the VOTC ROI (the paired samples Wilcoxon tests indicated a pFDR < 0.001 for Controls, Cataract-reversals, and Controls-Blurry2, and a pFDR < 0.005 for Controls-Blurry1). We also examined whether the Spearman’s correlation computed between the RDM extracted from the V1-like layer and the RDM extracted from the brain V1 ROI significantly differed between groups/conditions. Mann-Whitney independent samples t-tests showed a trend for a higher correlation in the Control group compared to all the others (Controls > Cataract-reversals, pFDR = 0.09; Controls > ConBlurry1, pFDR = 0.07; Controls > ConBlurry2, pFDR = 0.09). No other group differences emerged.

We repeated the same analyses also for the VOTC-like layer. The RDM extracted from the VOTC-like layer of the DNN exhibited a significant correlation with the RDMs extracted from the VOTC ROI in each group/condition (Spearman’s r: Controls 0.428, Cataract-reversals 0.411, Controls-Blurry1 0.371, and Controls-Blurry2 0.294; all p < 0.001 at the Wilcoxon one-sample test, FDR corrected). Additionally, as expected, within each group, this correlation value was significantly higher compared to the correlation between the RDM extracted from the same layer (VOTC-like layer) of the DNN and the RDM extracted from the V1 ROI (the paired samples Wilcoxon tests indicated a pFDR < 0.001 for Controls, Cataract-reversals, and Controls-Blurry1, and a pFDR < 0.05 for Controls-Blurry2). We also examined whether the Spearman’s correlation computed between the RDM extracted from the VOTC-like layer and the RDM extracted from the brain VOTC ROI significantly differed between groups/conditions. Mann-Whitney independent samples t-tests showed a trend for a higher correlation in the Control group compared to the ConBlurry1 (pFDR=0.06), a significant difference between the Control group and the ConBlurry2 condition (pFDR<0.001) and a significant difference between the Cataract-reversal group and the ConBlurry2 (pFDR<0.001).

In summary, these findings align with previous literature (Cichy et al., 2016) and suggest that earlier DNN layer intended to mimic V1 shows a stronger correlation with brain representations from V1 than from VOTC across all groups, while the later DNN layer intended to mimic VOTC exhibits a stronger correlation with brain representations from VOTC than from V1 in every group. Interestingly, we also observed more fine-grained results regarding group differences. Specifically, the correlation with the V1-like layer tends to be higher with the V1 representation in the Control group compared to the Cataract-reversal, ConBlurry1, and ConBlurry2 groups. In contrast, the correlation with the VOTC-like layer appears higher in the Control and Cataract groups compared to the ConBlurry1 and ConBlurry2 conditions. Although some of these results are not statistically significant, the trends clearly align with the direction of our brain experiment findings.

#### RSA analyses: correlation of RDMs extracted from different layers and conditions of DNN with representational models (HmaxC1 and categorical)

Here, we directly tested whether the two representational models selected to examine low-vs. high-level visual representations in the occipital ventral stream (i.e., the HmaxC1 and categorical models) would also apply to visual representations across different layers of the DNN.

These results are depicted in fig. 7C. We observed a significant correlation between the RDM extracted from the V1-like layer and the HmaxC1 model across all DNN conditions: trained and tested with typical images (pFDR < 0.001), trained and tested with filtered images (pFDR < 0.05), and trained with typical images and tested with filtered images (pFDR < 0.005). Moreover, the Spearman’s correlation in the Control-like DNN condition was significantly higher compared to the Cataract-reversals-like condition (pFDR < 0.05) and showed a trend toward significance compared to the ConBlurry-like condition (pFDR = 0.09).

Similarly, we observed a significant correlation between the RDM extracted from the VOTC-like layer and the categorical model across all DNN conditions: trained and tested with typical images (pFDR < 0.001), trained and tested with filtered images (pFDR < 0.001), and trained with typical images and tested with filtered images (pFDR < 0.005). Furthermore, the Spearman’s correlation in the Control-like DNN condition showed a trend toward significance compared to the ConBlurry-like condition (pFDR = 0.09).

In summary, these findings support our brain results obtained with RSA using the HmaxC1 and categorical models. They show that in V1-like layer, the cataract-like condition exhibits an impaired representation of low-level visual features compared to the control-like condition, resembling the ConBlurry-condition. In contrast, in VOTC-like layer, the cataract-like condition shows a preserved categorical representation like the control-like condition, while categorical representation is still impaired in the ConBlurry-like condition.

While some trends are not statistically significant, the numerical patterns observed in these DNN analysis closely replicate our brain data, reinforcing the direction of the results. Moreover, the following decoding analyses provide additional statistical supports to our findings.

#### Between- and within-category decoding analyses in different layers and conditions of the DNN

The fMRI data show reduced between-category decoding in the VOTC for ConBlurry 1 and 2 compared to both controls and cataract-reversal subjects, while within-category decoding is reduced in V1 for both ConBlurry 1 and 2 and cataract-reversal subjects compared to controls. With this final analysis, we sought to determine whether a similar pattern was also present in the DNN layers.

All decoding results across different DNN layers are presented in fig. 7D.

As shown in the left panel of fig. 7D, the permutation test revealed that the between-category decoding performance exceeded chance level (20%) in the VOTC-like layer across all DNN conditions (all pFDR < 0.001): trained and tested with typical images (Control-like condition, DA: 77%), trained and tested with filtered images (Cataract reversals-like condition, DA:94%), and trained with typical images and tested with filtered images (Controls blurry-like condition, DA:59%). Additionally, the permutation analysis indicated significant differences between all pairwise condition comparisons: decoding accuracy in the Cataract reversals-like condition was significantly higher than both the Control-like condition (pFDR < 0.001) and the Control blurry-like condition (pFDR < 0.001). Furthermore, decoding accuracy in the Control-like condition was significantly higher than in the Control blurry-like condition (pFDR < 0.001).

The results for the within-category decoding analyses are presented in fig. 7D in the right box. The permutation test revealed that within-category decoding performance exceeded chance level (16.7%) in the V1-like layer for all categories in the Control-like condition, where the DNN was trained and tested with typical images (Bodies: DA=37%, p=.003; Faces: DA=33%, p=.01; Houses: DA=90%, p=.0002; Tools: DA=60%, p=.0002; Words: DA=83%, p=.0002). However, in the cataract-like condition, where the DNN was trained and tested with filtered images, the within-category decoding accuracy never significantly exceeded chance level in any of the categories (Bodies: DA=10%, p=.89; Faces: DA=6%, p=.89; Houses: DA=20%, p=.89; Tools: DA=13%, p=.89; Words: DA=17%, p=.89). Finally, in the control blurry-like condition, where the DNN was trained with typical images and tested with filtered images, the within-category decoding accuracy was significantly higher than chance level for Houses (DA=33%, p=.02) and Words (DA=33%, p=.02), but not for the other three categories (Bodies: DA=17%, p=.49; Faces: DA=6%, p=.89; Tools: DA=20%, p=.38). Regarding the differences between conditions, the permutation tests revealed significant differences between the Control-like condition and the other two conditions in all categories (Bodies: Control-like vs Cataract-like p=0.03, Control-like vs ConBlurry-like p=0.02; Faces: p=0.005 for both comparisons; Houses, Tools, and Words: p=0.0002 for both comparisons). No difference between the Cataract-like and Control blurry-like conditions did emerge for any category.

These decoding analyses clearly show, as observed in the brain results, that in the V1-like layer, the Cataract-like condition exhibits impaired processing, behaving more like the ConBlurry-like condition than the Control-like condition. In contrast, when examining VOTC-like processing, the Cataract-like condition displays an intact categorical representation compared to the ConBlurry-like condition.

## Discussion

We investigated how a brief period of visual deprivation during the neonatal stage affects the development of the human object recognition system. Using a multivariate framework incorporating human brain imaging and artificial neural networks, we examined the impact of visual deprivation early in life on the development of the visual object recognition system. Our findings revealed a preserved categorical representation in the ventral visual pathway (VOTC) despite a brief and transient period of visual deprivation early in life and permanent alterations in the response properties of the primary visual cortex (V1) for low-level visual features.

The V1 of neonates demonstrates an advanced level of connectivity comparable to the one observed in adults (Burkhalter, 1993; Coogan & Van Essen, 1996; Horton & Hocking, 1997), exhibits ocular and orientation columnar organization (Blasdel et al., 1995; Crair et al., 1998) displays selectivity for visual features similar to adult levels (Wiesel & Hubel, 1974) and advanced retinotopic organization (Arcaro et al., 2017). Moreover, the eccentricity-related resting-state connectivity profile of early visual cortex (EVC) appears roughly preserved in congenitally and permanently blind adults (Bock et al., 2015; Striem-Amit et al., 2015). Do these findings suggest that EVC development occur independently of sensory experience? Our results demonstrate otherwise by revealing that cataract-reversal individuals have altered representation of low-level visual features, as measured by reduced RSA correlation with a computational V1-like model (fig. 5) and by reduced item-decoding analyses within each category (fig. 6). These results align with former animal studies indicating lasting functional deficits in V1 properties in kittens, monkeys, and mice exposed to early and transient visual deprivation (Crair et al., 1998; Knudsen, 2004; Wiesel, 1982). Studies involving human participants after a transient period of early visual deprivation also suggested lasting deficit in low- and mid-level visual abilities (Maurer et al., 2006; Maurer & Lewis, 2001; McKyton et al., 2015) and alteration in the response properties of EVC (Heitmann et al., 2023). Altogether, our results show how early visual experience plays a crucial role in maintaining and refining the anatomical and functional development of early visual circuits, despite a prototopographic organization at birth (Heitmann et al., 2023; Huberman et al., 2008).

In contrast to the notion that V1 displays a relatively high level of functional and structural maturity at birth (Blasdel et al., 1995; Coogan & Van Essen, 1996; Wiesel & Hubel, 1974), the extrastriate visual system has long been thought to display rather immature neonatal functional and anatomical organization (Bourne & Rosa, 2006; Kiorpes & Movshon, 2003; Spriet et al., 2022; Zhang et al., 2005). For instance, categorical selectivity in VOTC shows a protracted period of maturation that extends into adolescence and sometimes even adulthood as for faces (Golarai et al., 2015; Nordt et al., 2021). Combined with the idea that downstream higher-level regions acquire their functional tuning based on structuring inputs they receive from earlier regions in the cortical hierarchy, this led to the assumption that visual deprivation induces a cascaded pattern of impairments, with its magnitude increasing as a function of the synaptic distance from the earlier visual input (Hyvärinen et al., 1981; Maurer et al., 2007). Recent studies on human cataract-reversal individuals suggested greater alterations of visual processing implemented in extrastriate compared to striate brain regions (Pitchaimuthu et al., 2021; Sourav et al., 2018, 2020), suggesting that the functional impact of visual deprivation becomes more prominent downstream in the visual processing hierarchy (Röder & Kekunnaya, 2022). A preliminary report even suggested that monkeys raised in darkness for a year show no impairment in V1 but impairment in VOTC, where activity for visual stimuli is observable but not selective for specific categories (M. Arcaro et al., 2018). Our findings challenge these assumptions. Despite clear differences in V1 response properties, we did not observe differences between the cataract-reversal group and control group in VOTC response. In this region, the reliability of activity patterns for each item was equivalent between the two groups (split-half correlation measurements; see fig. 4), and we observed a comparable categorical representation expressed by univariate contrasts (fig. 3), RSA correlation with a categorical model (fig. 5), and multiclass decoding analyses between categories (fig. SI 2). Such convergence in results using multiple analytical streams, clearly challenges the view that visual areas most mature at birth are the most resilient to experience-dependent plasticity (M. J. Arcaro & Livingstone, 2017; Röder & Kekunnaya, 2021) and interrogate the notion that categorical domain formation is primarily influenced by early visual experience (M. J. Arcaro & Livingstone, 2021). Our study instead suggests that early postnatal visual experience is not a prerequisite for developing categorical selectivity in VOTC, in agreement with the fact that cataract-reversal individuals have no known deficits in visual categorization (Maurer et al., 2005), including discriminating faces from non-faces (Geldart et al., 2002).

An important difference between our and previous studies is the length of visual deprivation. We examined the impact of a brief period of postnatal visual deprivation, lasting approximately 100 days on average. By contrast, previous studies examined significantly longer periods of deprivation in both humans (Heitmann et al., 2023; Sourav et al., 2018) and primates (M. Arcaro et al., 2018; Hyvärinen et al., 1981). Crucially, the time between the eyes opening and the test in previous studies was significantly shorter than the several years that elapsed between surgery and testing in our study (Table 1). In our study we tested cataract-reversal individuals many years after surgery. This extensive visual experience might have contributed to the maintained categorical representation in VOTC. Indeed, previous behavioral studies have documented a gradual recovery of many visual abilities following congenital cataract surgery. For example, visual acuity in children treated early continues to improve until about age two, even if it doesn’t reach typical levels (Lewis et al., 1995; Maurer & Lewis, 2001). Longitudinal studies of individuals treated after extended blindness also show that, despite some low-level impairments (e.g., reduced acuity), these individuals demonstrate proficiency in many mid- and high-level visual tasks, including shape matching, visual memory, image segmentation, and face discrimination, localization & identification, and gender classification (Ostrovsky et al., 2006; Sinha & Held, 2012). This behavioral recovery suggests parallel recovery at the neural level, particularly in brain regions involved in these visual functions, which would include VOTC. Longitudinal studies examining brain changes in various regions of the visual cortex following cataract removal are essential to understand the mechanisms supporting this recovery trajectory at the brain level.

Importantly, previous studies involving cataract reversal participants with similar period of transient deprivation reported behavioral impairments in fine-grained categorical processing, mainly in facial recognition, in cataract-reversal subjects (De Heering & Maurer, 2014; Le Grand et al., 2001; Robbins et al., 2010). These impairments may stem from initial visual analysis deficits in V1 or potential impairments in VOTC representation at more detailed levels. We do not rule out the possibility of VOTC impairments in more fine-grained intra-categorical representations, in particular in tasks that require feature binding (McKyton et al., 2015; Putzar et al., 2007). For the face domain specifically, previous research has indeed suggested that the neural processes involved in discerning fine details of a specific face may vary from those implicated in recognizing generalized facial configurations (Kobylkov & Vallortigara, 2024; Tsao & Livingstone, 2008) and holistic face processing, a hallmark of identity recognition, has been shown to be altered in cataract-reversal patients despite intact face discrimination (Geldart et al., 2002; Putzar et al., 2010).

Our results support the existence of varying sensitive periods along the visual processing hierarchy, leading to distinct developmental impacts of sensory deprivation (Maurer, 2017; Maurer & Lewis, 2013; Röder et al., 2021). This aligns with the idea that different visual skills—and the brain regions supporting them—may vary in their susceptibility to early visual deprivation (Sinha & Held, 2012) supporting the existence of multiple sensitive periods across different brain regions (Maurer, 2017; Maurer & Lewis, 2013; Röder et al., 2021). However, which brain regions are more or less affected by early visual deprivation remains controversial. In striking contrast with the idea that impairments increases from V1 to more downstream regions (Röder et al., 2021), we instead found that early blindness compromises the representation of low-level visual information in V1, while visual categorical representation in VOTC appears unaffected.

To explore whether V1 impairment was due to early visual deprivation or current visual impairment at the time of testing, we included several control analyses. First, we extracted and regressed out information on each subject’s eye movements during the experiment (fig. 2C) to minimize data variance due to different eye movement profiles across groups. Correlation analyses between brain data and visual acuity values for the cataract-reversal individuals was not significant (e.g., fig. 4H, fig. 5E), suggesting observed V1 results were not linked to visual acuity at the moment of testing. Additionally, we included two control conditions where participants with normal vision underwent the experiment with degraded visual presentation, mimicking cataract-reversal individuals’ vision. In this control group, we observed that degraded visual input produced widespread impairment across both V1 and VOTC, indicating that typical visual development involves cascading effects from V1 to VOTC when visual input quality is reduced. However, cataract-reversal individuals showed a distinctly different profile: their impairment was isolated to V1, with categorical representation in VOTC remaining intact.

In addition, we investigated whether hierarchical visual processing as implemented in artificial neural network could serve as a relevant model to simulate the functional consequence of an early and transient period of visual deprivation. As expected, early DNN layer representation correlated more with cortical representations in V1, and late DNN layer representation correlated more with the anterior portion of the ventral visual pathway (Cichy et al., 2016). In line with our results, the V1-like layer representation was higher with the brain V1 representation of the control group compared to cataract-reversals and visually-degraded conditions, while the VOTC-like layer representation was similar in controls and cataract-reversals but lower in visually-degraded conditions (fig. 7B).

Building upon this similarity, we independently manipulated certain aspects of the network architecture and the training-testing dataset to mimic the three groups included in our fMRI experiment: Control-like, Cataracts-like, and ConBlurry-like conditions (fig. 7A). We then replicated with the DNN data the RSA and the decoding analyses that we previously performed on the brain data. Our DNN results closely match the brain results. Both the RSA analysis with the low-level visual model and the categorical model (fig. 7C) and the decoding analyses (fig. 7D) pointed to an impairment in the V1-like layer for the processing of low-level visual properties of the stimuli both in the Cataracts-like and ConBlurr-like conditions compared to the Control-like condition. Instead, the categorical representation in VOTC-like layer was altered only in the Con-Blur like condition compared to both the Control-like condition and the Cataract-like condition. These findings in artificial neural networks therefore support our biological data, revealing a distinction between living with reduced visual acuity and perceiving degraded visual stimuli. Our control experiment and DNN results demonstrate that in typical visual development, degraded visual input leads to cascading impairment in V1 and VOTC.

Our data suggest that the brain employs alternative mechanisms to preserve categorical representation and selectivity in VOTC, distinguishing it from the acute effects of degraded vision. Here the DNN results provide a unique insight and help to interpret the neural data. More in general, these results also reveal that DNNs represent an interesting approach to evaluate the consequence of an early and transient period of deprivation and therefore pave the way to manipulate various features of deprivation that are notoriously difficult to control in humans (e.g., exact period and severity of deprivation) to generate new hypotheses on the functional consequences of sensory deprivation (Vogelsang et al., 2018).

The maintenance of categorical selectivity in VOTC despite early visual deprivation and V1 functional impairment could potentially be related to its connectivity with downstream regions that could exert a top-down regulation for the development of the categorical coding of VOTC (Howell et al., 2020; Li et al., 2020; Mattioni et al., 2020, 2022; Osher et al., 2016; Saygin et al., 2012, 2016). Categorical domains within the VOTC might have an inherent predisposition to process these specific categories such as faces, words, landscapes through specialized connections to temporo-parieto-frontal networks related to social cognition, linguistic processing, and spatial interaction with the environment (Hannagan et al., 2015; Mahon & Caramazza, 2011; Powell et al., 2018). Indeed, besides enabling object recognition, VOTC is likely a foundational brain region in determining the relevance of objects to behavior, for instance to engage in social interaction, object manipulation and in navigating the environment (Conway, 2018). Such pattern of large-scale connectivity might explain in part the maintained categorical coding of VOTC despite deprivation and EVC alteration (Powell et al., 2018).

Early visual deprivation triggers crossmodal activity for touch or sounds in EVC (Bavelier & Neville, 2002; Frasnelli et al., 2011; Van Ackeren et al., 2018), even years after visual restoration (Collignon et al., 2015; Dormal et al., 2015; Guerreiro et al., 2016), and sound stimulation can suppress early visual responses in cataract reversal individuals (Sourav et al., 2024). Crossmodal responses during a sensitive period could occupy synaptic space in the EVC, potentially interfering with visual recovery. In VOTC, crossmodal recruitment by sound in blind individuals appears to follow the category-selective profile of the region and is also observable in sighted individuals (Amedi et al., 2001; Mattioni et al., 2020; Pietrini et al., 2004; Van Den Hurk et al., 2017), suggesting that it may promote functional maintenance (Heimler et al., 2014; Makin & Krakauer, 2023), rather than interfere with sight recovery. In other words, the selective alteration of visual function in specific regions might also relate to region-specific impact of crossmodal reorganization triggered by early blindness (Heimler et al., 2014).

By showing how a brief period of early-life visual deprivation permanently affects information encoding in EVC while leaving the categorical coding in VOTC intact, our study highlights how distinct regions in the human object visual recognition system are differently affected by a transient period of postnatal blindness. Our results challenge the conventional belief that high-level brain regions predominantly depend on early visual experiences and intact downstream visual input for their development, and instead show how the development of the categorical coding in VOTC shows resilience to deprivation and altered downstream input. Finally, our study offers insights into how the brain processes visual information in the absence of typical visual experiences, paving the way to focused rehabilitative approaches based on the patient’s selective deficits as observed in our study.

## Acknowledgements

We would like to express our gratitude to Sally Stafford and Joy Williams who have helped with recruiting participants and the data acquisition, respectively. Computational resources have been provided by the supercomputing facilities of the Université catholique de Louvain (CISM/UCL) and the Consortium des Équipements de Calcul Intensif en Fédération Wallonie Bruxelles (CÉCI) funded by the Fond de la Recherche Scientifique de Belgique (F.R.S.-FNRS) under convention 2.5020.11 and by the Walloon Region. OC is a senior research associate at the Fond National de la Recherche Scientifique de Belgique (FRS-FNRS).

## Author Contributions

Conception and design: SM, MR, XG, DM, VG, AC, HoB, OC. Administrative support: SM, MR, XG, DM, TL, OC; Provision of study materials or patients: SM, MR, XG, DM, TL, OC; Collected data: SM, MR, XG, JN, ZL, OC. Analyzed data: SM, AC. Wrote the original draft: SM, OC. Writing - Review & Editing: MR, XG, DM, VG, AC, HoB. Main funding acquisition: OC.

## Data availability statement

Raw f/MRI data are not publicly available as full anonymity of the participants cannot be guaranteed, even after defacing the MRI images and due to the lack of explicit consent from our participants. Preprocessed data, brain masks and statistical outputs can be found on OSF: DOI 10.17605/<u>OSF.IO/BECDR</u>. Codes used to process the data can be found on GitHub: https://github.com/SteMat9787/Neurocat3.

