## Supplemental Material for "Impact of a transient neonatal visual deprivation on the development of the ventral occipito-temporal cortex in humans"

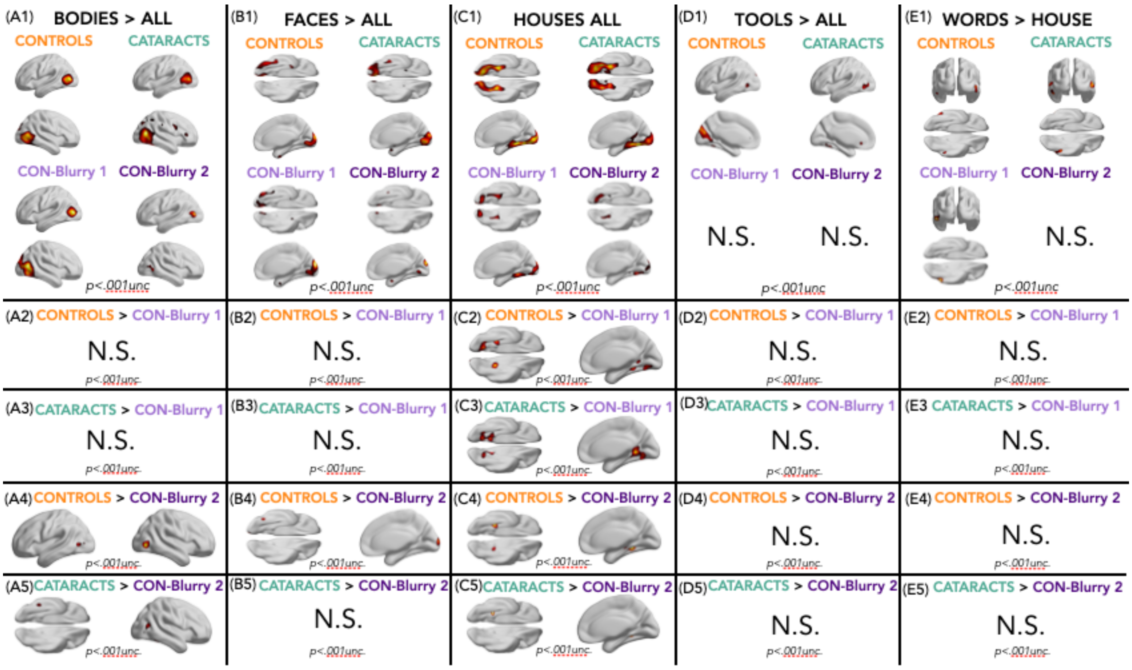


**SI Figure 1: Results from Univariate Analysis across category.** Each column displays univariate analysis results for one category: bodies **(Panels A1–A5)**, faces **(Panels B1–B5)**, houses **(Panels C1–C5)**, tools **(Panels D1–D5)**, and words **(Panels E1–E5)**. The top row shows univariate results for each category within each group and condition, with brain maps thresholded at p < 0.001 unc. For each category, the analysis contrasts that category against all others, except for words **(Panel E1)**, where we report the contrast words > houses, as words > all did not yield significant results at p < 0.001 unc. across groups. Rows 2 through 5 show groups' contrasts for categories with at least one significant difference between groups at p < 0.001 unc. Groups' contrasts without significant findings for any category are omitted. BrainNet Viewer was used for the visualization of brain maps (Xia et al., 2013).

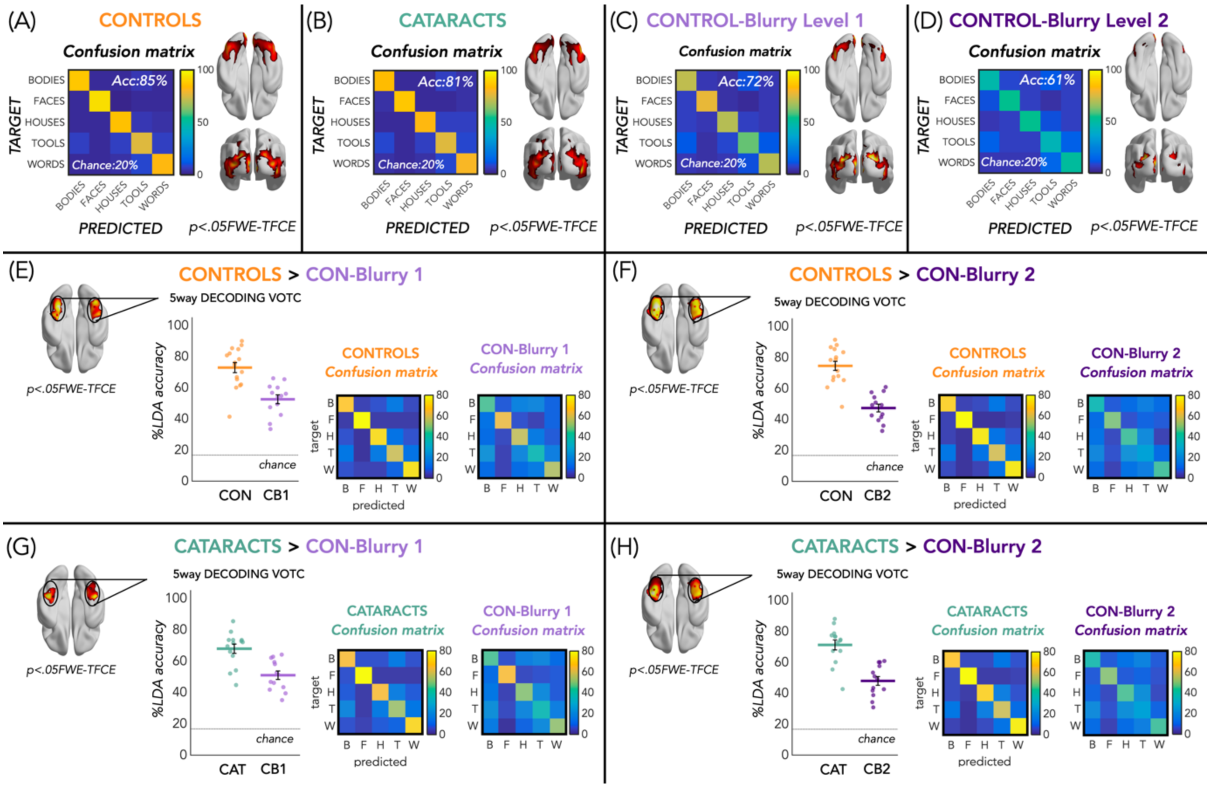


**Supplemental Figure 2. Results from Searchlight 5-way decoding analysis.** The top row presents results from the searchlight analysis within each group and condition: controls (**Panel A**), cataract-reversals (**Panel B**), Control-blurry exp 1 (**Panel C**), and Controls-blurry exp 2 (**Panel D**). Within each panel, the brain maps with regions showing significant decoding accuracy for the 5 categories are depicted on the right, and the confusion matrix extracted from these brain regions are represented on the left. The second row shows contrasts between the Control group and the Con-Blurry1 in **Panel E**, and Control group and the Con-Blurry2 in **Panel F**. The third row shows contrasts between the Cataract-reversal group and the Con-Blurry1 in **Panel G**, and Cataract-reversal group and the Con-Blurry2 in **Panel H**. All these maps are reported with a $p < 0.05$ FWE and Threshold-Free Cluster Enhancement (TFCE) corrected threshold. For regions showing group differences, confusion matrices are represented from both groups, and decoding accuracies are reported as dot plots for visualization purposes only. BrainNet Viewer was used for the visualization of brain maps (Xia et al., 2013).

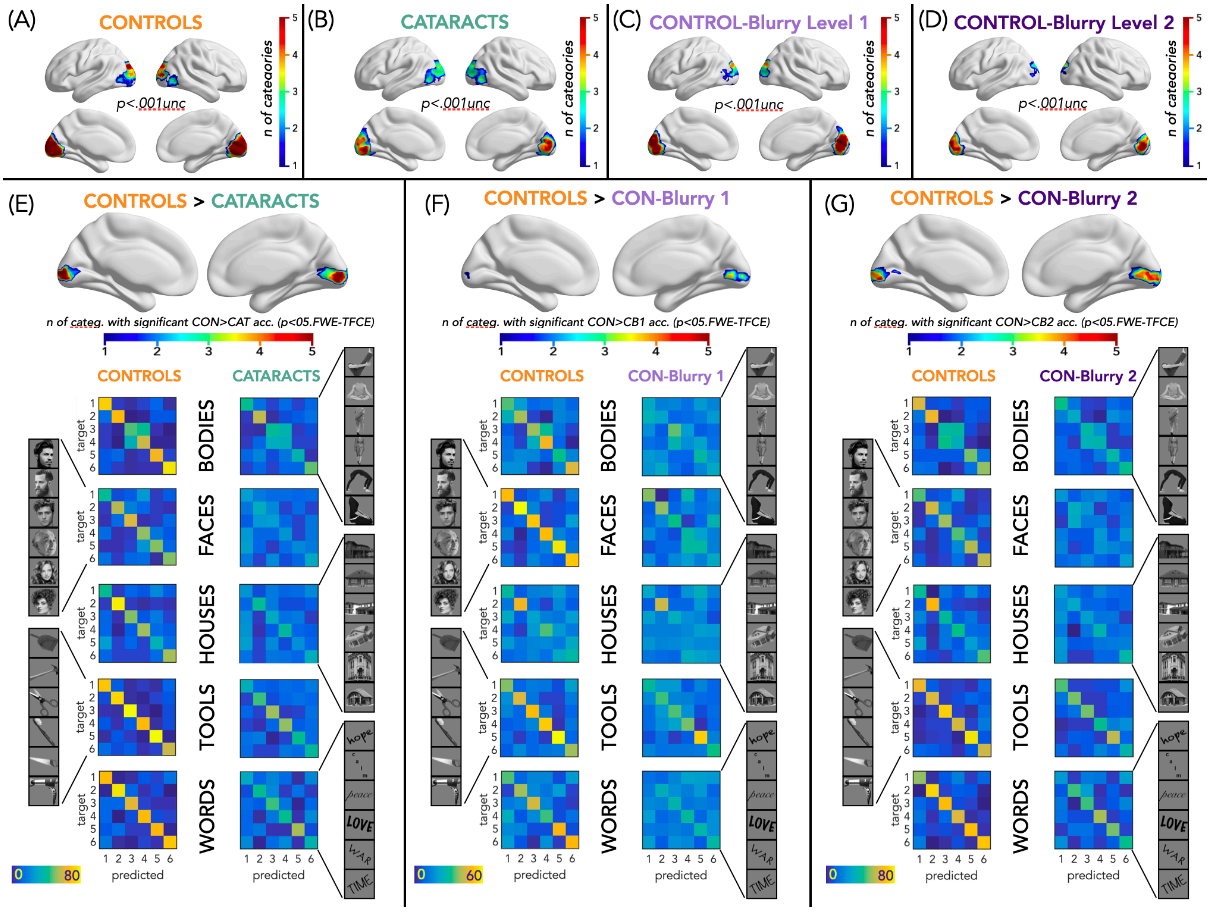


**Supplemental Figure 3. Within-category decoding analyses: overlap between categories.** These results depict the overlap of the searchlight results for the five different decoding analyses, each conducted on a subset of the data, focusing on one category at a time. The top row presents the results from the searchlight analysis within each group and condition: controls (**Panel A),** cataract-reversals (**Panel B**), control-blurry level 1 (Panel C), and controls-blurry level 2 (**Panel D**). Within each panel, we display the overlap of the five brain maps (one for each within-category decoding analysis) with regions exhibiting significant decoding. Regions in dark blue signify significant decoding accuracy for only one category, light blue for two categories, green for three categories, orange for four categories, and red for all five categories. Brain maps are depicted with a threshold at $p < 0.001$ uncorrected. The second row illustrates contrasts between the Control group and the other three groups: Control > Cataract in **Panel E**, Control > Con-Blurry1 in **Panel F**, and Control > Con-Blurry2 in **Panel G**. Similarly, these represent the overlap of the five brain maps, with colors ranging from blue to red indicating the number of categories. For example, brain regions in red denote areas where within-category classification accuracy was higher in the controls compared to the other group across all five categories. These maps are reported with a threshold corrected for Family-Wise Error (FWE) and Threshold-Free Cluster Enhancement (TFCE) at $p < 0.05$ within a visual mask. For regions displaying group differences, confusion matrices are presented for both groups and for each category individually. BrainNet Viewer was used for the visualization of brain maps (Xia et al., 2013).
